# An area-resolved phylogeography of bulbous barley (*Hordeum bulbosum*; Poaceae)

**DOI:** 10.1101/2025.01.20.633865

**Authors:** Frank R. Blattner, Simon Pfanzelt, Christina Koch, Axel Himmelbach, Agostina B. Sassone, Abdulnaser Kh. Abdulkarim, Lulëzim Shuka, Dörte Harpke

## Abstract

*Hordeum bulbosum*, the closest relative of barley (*H. vulgare*), is an important source of resistance genes for cereals in the Triticeae. This perennial and mainly outcrossing species occurs with two cytotypes: diploids thrive in the western and central parts of the Mediterranean, while autotetraploids extend from Greece mainly eastwards to western Asia. To elucidate *H. bulbosum*’s origin, colonization patterns, population relationships, and distinctions between the two cytotypes, we determined ploidy and performed genotyping-by-sequencing analyses on 314 individuals from across the species’ distribution range. Our results revealed two distinct lineages within diploid *H. bulbosum*: individuals from Libya *vs*. all other diploids. Southeastern Italian populations were the origin for the species expansion eastward into Albania/Greece and westward into mainland Italy, Sicily/Sardinia, Tunisia, and Spain. The tetraploid cytotype originated early in the evolution of the species, thus retaining alleles found in extant Libyan diploids. Tetraploids underwent local introgression from diploids in Greece, where also a secondary origin of tetraploids was detected. We found that ecoclimatic conditions alone cannot account for the clear geographic separation of the cytotypes, as habitats across much of the central Mediterranean would be suitable for both. We conclude that minority cytotype exclusion is the most plausible explanation for the distinct distribution patterns observed. While both cytotypes are morphologically indistinguishable, diploid populations typically occur in scattered stands within their range, while tetraploids tend to form larger, often contiguous populations.

## Introduction

Bulbous barley (*Hordeum bulbosum* L.) is a grass native to open habitats from Portugal and Morocco in the west to Tajikistan in the east (Jørgensen, 1982). Although generally quite similar to barley (*Hordum vulgare* L.), the conspicuous bulbous structures at the base of its stems make these species easily distinguishable. In contrast to barley, *H. bulbosum* is perennial, predominantly outcrossing, and occurs with an autotetraploid in addition to the diploid cytotype. Both cytotypes are strongly separated geographically, with the diploid occurring in the central and western Mediterranean, and the tetraploid occurring from central Greece eastward (Katznelson & Zohary, 1967; Jørgensen, 1982). Bulbous barley diverged from the barley lineage approximately 4.5 million years ago (Mya; Brassac & Blattner, 2015) and is the only species in the secondary gene pool of barley (Bothmer *et al*., 1995; Blattner, 2009, 2018), allowing crossbreeding, albeit with reduced offspring fertility (Subramanyam & Bothmer, 1987). Bulbous barley has long been used to easily produce haploid barley lines through crosses between the two species (Jensen, 1977) and was recognized early on as a potential source of resistance factors not present in the crop (Konzak *et al*., 1951; Jones & Pickering, 1978; Pohler & Szigat, 1982; Walther *et al*., 2000; Johnston *et al*., 2009; Feng *et al*., 2024). Genomic tools developed in the last decade (Wendler *et al*., 2014, 2015; Feng *et al*., 2024) will soon support targeted gene transfer from *H. bulbosum* to barley and other cereals, and also foster approaches to domesticate this species as a perennial grain crop (Westerbergh *et al*., 2018; Chapman *et al*., 2022; Fuerst *et al*., 2023). Despite the importance of *H. bulbosum* for future crop improvement, a systematic assessment of the genetic structure of its wild populations is still lacking.

To address this knowledge gap, we sampled a representative set of *H. bulbosum* populations from across its range, determined the ploidy levels of individuals, constructed habitat suitability maps, quantified the ecoclimatic niche overlap of di- and tetraploid cytotypes, and performed a phylogeographic analysis based on genome-wide DNA polymorphisms obtained through genotyping-by-sequencing (GBS; Elshire *et al*., 2011), a reduced genomic representation sequencing method. With these analyses, we aim to (i) determine the area of origin of the species and its tetraploid cytotype, (ii) infer colonization routes, (iii) identify gene flow between populations and cytotypes, and (iv) try to identify ecological differences that led to the strong geographic separation between di- and autotetraploid populations.

## Materials and Methods

### Plant materials

*Hordeum bulbosum* populations in the central and western Mediterranean (Fig. 1) were visited between end of May and early October. About 10 ripe grains for each of three to seven individuals per population (depending on population size) were collected, bagged individually per mother plant, and transported to the Leibniz Institute of Plant Genetics and Crop Plant Research (IPK Gatersleben, Germany). Seeds were germinated in pots and one seedling per mother plant and two to five seedlings per population were further grown, regularly repotted and kept till flowering stage for leaf and seed harvesting. The population samples were complemented by accessions from the germplasm collections of the genebanks of the IPK Gatersleben and US Department of Agriculture (USDA National Small Grains Collection) to also represent genotypes from northern Africa and tetraploids from western Asia. Collecting and exchange of seeds followed the regulations of the respective countries and genebanks. One herbarium voucher per analyzed population was deposited in the herbarium of the IPK Gatersleben (GAT). The entire *H. bulbosum* dataset consisted of 230 diploid individuals from seven countries and 84 individuals representing the tetraploid lineages from nine countries. As barley is sister to bulbous barley (Brassac & Blattner, 2015), five barley accessions, i.e. *Hordeum vulgare* subspp. *vulgare* and *spontaneum* (K.Koch) ASCH. & GRAEBN., were included as outgroup. For details of the analyzed populations see Supplementary Online Table S1.

**Figure 1.**
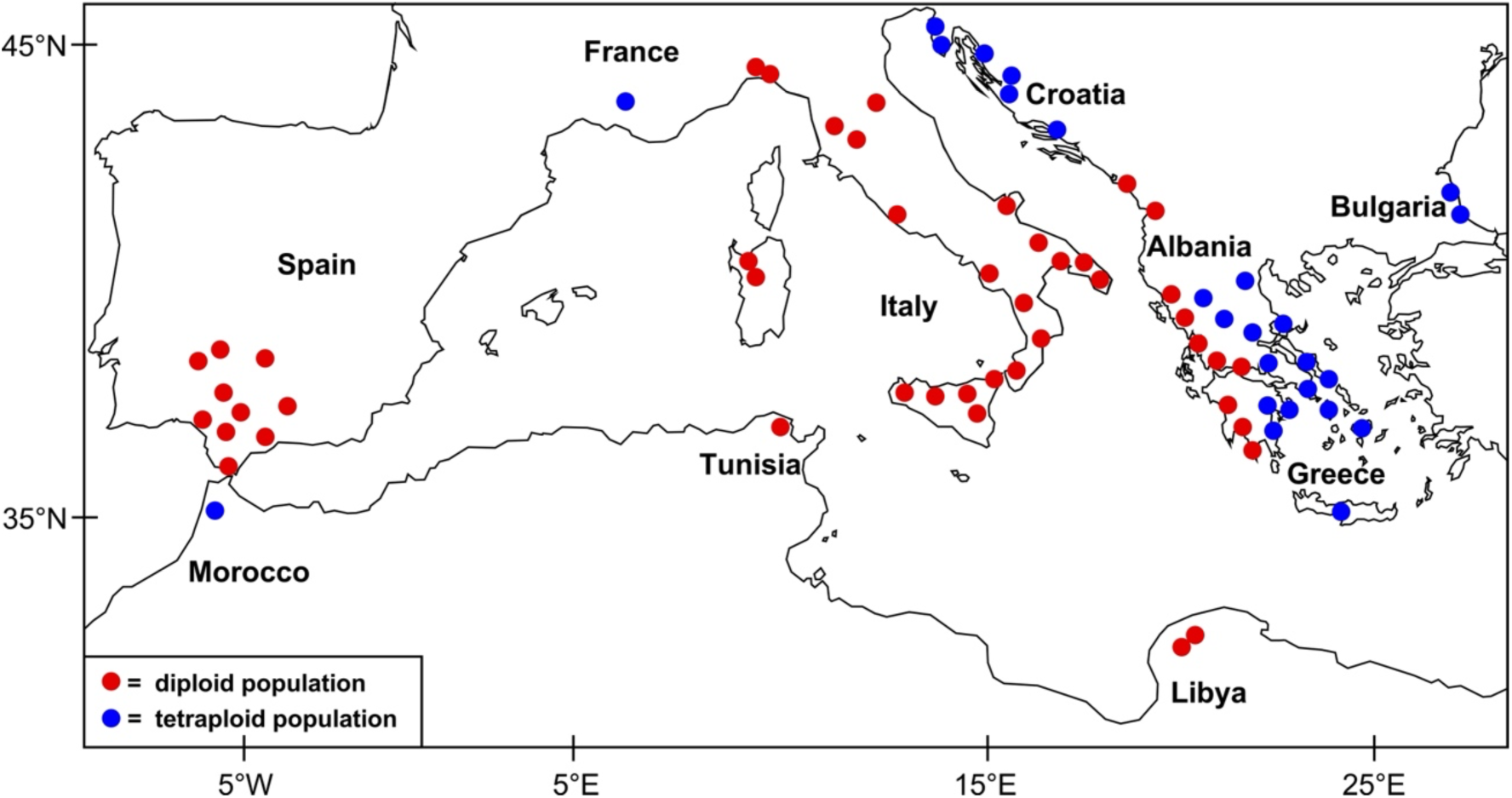
Collection sites for *H. bulbosum* in the central and western parts of the Mediterranean basin, indicating the borders between di- and tetraploid populations. For details of the Asian tetraploids included in the analyses see Supplementary Table S1.

### Ploidy determination

Genome sizes of all individuals were measured using fresh leaves and propidium iodide (PI) as dye with a Cyflow Space (Sysmex Partec) flow cytometer, essentially following the procedure described by Jakob *et al*. (2006). We used *Vicia faba* L. cv. ‘Tinova’ (IPK genebank accession FAB 602; https://doi.org/10.25642/IPK/GBIS/33373) with a 2C genome size of 26.21 pg as size standard, as the expected di- and tetraploid 2C values of *H. bulbosum* should be around 9 pg and 17 pg, respectively (Jakob *et al*., 2006).

### Molecular methods

DNA extraction was conducted with DNeasy Plant Mini kits (Qiagen) from young leaves dried in silica gel. DNA extraction followed the protocol of the manufacturer. DNA quality was checked on agarose gels and for quantification the Qubit dsDNA High Sensitivity assay kit (Thermo Fisher Scientific) was used. To obtain genome-wide SNP data, we implemented a two enzyme GBS method (Poland *et al*., 2012), utilizing the restriction enzymes *Pst*I and *Msp*I. Library preparation, individual barcoding and sequencing were conducted as described in Ala *et al*. (2025).

### Data preparation

Barcoded reads from the 319 samples were de-multiplexed using the CASAVA pipeline 1.8 (Illumina). Adapter trimming of GBS sequence reads was conducted with CUTADAPT (Martin, 2011) within IPYRAD v.0.9.87 (Eaton & Overcast, 2020). GBS reads were clustered in IPYRAD. We set the minimal number of samples to possess a certain locus to 90% of the individuals and the clustering threshold of reads within and between individuals to 0.90. The default settings were used for the other parameters except for the maximum depth (1000). After running the IPYRAD pipeline, three datasets were generated in the last step of IPYRAD, i.e. (i) one with all di- and tetraploid individuals including also *H. vulgare*, (ii) excluding the latter and restricting the dataset to *H. bulbosum* individuals, and (iii) one with only diploid *H. bulbosum* individuals. For GBS-based phylogenetic analyses the first dataset was modified by including or excluding *H. vulgare* and/or di- or tetraploid individuals. For population assignment analyses and principal component analysis (PCA) we used the datasets restricted to *H. bulbosum*.

### Population genetic analyses

The variant calling format (vcf) file generated by the IPYRAD pipeline of the *H. bulbosum* data sets underwent filtering using VCFTOOLS 0.1.16 (Danecek *et al*., 2011). This involved removing single nucleotide polymorphisms (SNPs) with a minor allele frequency below 0.05, variants with a depth below 10 or above 300, as well as indels. Only sites present in at least 90% of the sequences for the diploid *H. bulbosum* dataset and 70% for the di-plus tetraploid dataset were retained. The resulting vcf file was converted into STRUCTURE format using PGDSPIDER v 2.1.1.5 (Lischer & Excoffier, 2012). Model-based Bayesian population assignment analyses were conducted for single SNPs per fragment based on the filtered vcf files using the R package ‘LEA’ (Frichot & François, 2015) as well as STRUCTURE v2.3.4 (Pritchard *et al*., 2000). In LEA, population assignment was performed for K = 2–20 with 30 repetitions each, setting ploidy to two. The optimal K was determined based on the lowest cross entropy. Within STRUCTURE, a series of K = 2–20 was tested in ten independent runs, adopting the admixture model with correlated allele frequencies. The Monte Carlo Markov Chain (MCMC) was run for 10^6^ steps, discarding the initial 50% as burn-in. The Q-matrices from ten runs were averaged using CLUMPP v.1.1.2b (Mattias *et al*., 2007) applying 100,000 repetitions to calculate the mean of the ten iterations. STRUCTURE HARVESTER v0.6.94 (Earl *et al*., 2012) was employed, utilizing the Evanno method, to evaluate likelihood values across ten iterations for each of the K clusters, with the K exhibiting the highest ΔK considered as optimal.

The Q-matrices obtained from LEA and STRUCTURE at K = 5 for the diploid *H. bulbosum* dataset and K = 11 for the di-plus tetraploid dataset were organized using the ’tidyverse’ R package (Wickham & Wickham, 2019) and visualized with ’ggplot2’ (Wickham, 2016), depicting distinct ancestral clusters through color-coding. PCA results were computed in LEA and plotted with ggplot2. Individual pie-charts reflecting ancestral population assignments were obtained using the ’plotrix’ R package (Lemon, 2016).

### Phylogenetic analyses

To infer the phylogenetic relationships of *H. bulbosum* populations based on the GBS data, seven maximum parsimony (MP) analyses were conducted in PAUP* 4.0a169 (Swofford, 2002), with initially including (i) all 319 individuals and defining *H. vulgare* as outgroup, followed by analyses of the dataset including (ii) diploid *H. bulbosum* together with *H. vulgare* and (iii) only diploid *H. bulbosum*. In the latter analysis the Libyan individuals were defined as outgroup according to the results of the analyses before. As phylogenetic relationships among the tetraploid individuals appeared sensitive to the used outgroup, a separate set of analyses was run for tetraploid *H. bulbosum* individuals, including (iv) only tetraploids, and (v) *H. vulgare*, (vi) Libyan diploids, or (vii) Greek diploids as outgroups. For the dataset including only diploid individuals also a neighbor-joining (NJ) analysis was conducted in PAUP* to illustrate branch-lengths differences. Here we used the general time reversible (GTR) model of sequence evolution to calculate pairwise distances.

We conducted in all cases two-step MP analyses (Blattner, 2004) with an initial heuristic search with 1000 random-addition sequences (RAS) restricting the number of stored trees to 25 per repetition, tree-bisection reconnection (TBR) branch swapping, and steepest descent not enforced. The resulting trees were then used as starting trees in a heuristic analysis without restricting the number of sampled trees. Clade support was evaluated by bootstrap re-sampling. For the two large datasets including all diploids and di-plus tetraploids we used the fast-stepwise algorithm with 50,000 bootstrap re-sampling steps. For all other datasets we ran 500 bootstrap replicates with heuristic search settings as before except that we excluded the RAS step and used the simple-addition sequence. As fast-stepwise bootstrapping and excluding the RAS step results normally not in shortest trees in large GBS analyses, all MP bootstrap values we report here are rather conservative support estimations.

SVDQuartets (SVDQ) were calculated for the dataset consisting of diploid *H. bulbosum* together with *H. vulgare* individuals as outgroup in PAUP*. Individuals were partitioned (i) according to their geographic regions in 12 groups: Albania/S Croatia, Greece-Peloponnese, Greece-west, Italy-Apulia, Italy-Calabria, Italy-mainland, Italy-Sardinia, Italy-Sicily, Libya, Spain, Tunisia and outgroup, and (ii) by defining populations. The latter approach resulted in 19 groups with a finer subdivision of the Italian individuals. For Spain we defined neither regions nor populations, as none of the earlier analyses provided indications for phylogeographic structuring within the Iberian Peninsula. In SVDQ analyses we evaluated 25 million quartets (42% of all quartets), and trees were selected using Quartet Fiduccia and Mattheyses (QFM) assembly and the multi-species coalescent (MSC) as the tree model. For bootstrap support values we used 500 re-samples with the same settings as before but evaluating 2.5 million quartets only.

For the dataset including diploid *H. bulbosum* accessions together with *H. vulgare* as outgroup a maximum likelihood (ML) analysis applying the GTRGAMMA model and 100 parsimony starting trees in IQ-TREE v2.2.6 (Minh *et al*., 2020) was conducted. Statistical support was assessed via rapid bootstrapping with 500 replicates.

### Ecoclimatic niche models

To obtain a map of habitat suitability and model the ecoclimatic niche of di- and tetraploid *H. bulbosum*, a maximum entropy approach was employed, as implemented in MAXENT 3.3.3k (Philips *et al*., 2006). Occurrence records were based on sampling localities of plant material used for the phylogeographic analyses. Current bioclimatic data were downloaded from the WorldClim database v2.1 (climate data for 1970-2000; Fick & Hijmans, 2017). To avoid collinearity among bioclimatic layers, five layers were selected using hierarchical clustering. The used bioclimatic layers were mean diurnal range (the mean of the difference of maximum and minimum monthly temperatures), temperature seasonality, annual precipitation, as well as June values (corresponding to flowering time) of solar radiation and water vapor pressure. Pairwise Pearson’s correlation coefficients between the selected bioclimatic layers were <0.68. The ‘ENMeval’ R package (Muscarella *et al*., 2014) was used to build and compare several ecoclimatic niche models across a range of different settings. Ten thousand background points were drawn randomly from the study region, which was set to 10°W– 75°E and 27°N–47°N, covering the distribution range of *H. bulbosum*. Regularization values were 0.5, 1, 2, 3, 4; the sets of tested feature classes were L, LQ, H, LQH, LQHP, LQHPT; test data were partitioned using the “checkerboard1” setting, and the number of kfolds for evaluation was set to 4. The best model was selected based on ΔAICc values. Niche overlap was calculated with the R package ‘phyloclim’ (https://github.com/heibl/phyloclim/), using the statistics D (Schoener, 1968) and I (Warren *et al*., 2008).

## Results

### General properties of the dataset

Initially, a single dataset was created for all GBS-derived phylogenetic analyses in IPYRAD, including all 319 individuals. This dataset yielded 10,955 loci, resulting in a concatenated alignment of 1,241,989 bp (with 17.8% missing sites). From this dataset, two subsets were derived: (i) including only diploid accessions plus *H. vulgare* as outgroup, achieved by excluding all polyploid individuals, and (ii) including only tetraploids plus *H. vulgare*, along with Libyan and Greek populations as outgroups. These subsets were utilized to explore the influence of different outgroups on the topology of the tetraploids’ phylogenetic trees, either by excluding all outgroup individuals or using only one type of diploid as outgroup.

The dataset for population assignment analyses of diploid *H. bulbosum* individuals consisted of 230 samples from IPYRAD, comprising 11,587 loci and a SNP matrix of 99,346 SNPs with 10.75% missing sites. After filtering, 5,121 out of 99,346 possible sites were retained. The dataset for diploid and tetraploid individuals contained 314 samples, 10,567 loci, and a SNP matrix of 12,811 SNPs with 14.61% missing sites. Following filtering, 11,099 sites were retained.

### Ploidy level determination

Flow cytometry analysis revealed an average 2C genome size of 8.72 (SD ±0.29) pg for diploids and 16.81 (SD ±0.36) pg for tetraploids. Interestingly, within cytotype groups, no distinct differences in genome sizes correlating with geographic origins were observed. Triploids were absent in our dataset. However, one individual from the small Tribanj population in Croatia (FB21-004) exhibited a 2C genome size of 20.94 pg, indicating a pentaploid genome.

### Phylogenetic analyses

An initial phylogenetic analysis showed that the GBS reactions of replicated individuals were highly reproducible, resulting in clades where all replicates clustered together separate from other closely related individuals. For all further analyses only a single dataset of the replicated individuals was included.

Out of the 1,241,989 bp in our alignment for the initial analysis including all 319 *H. bulbosum* and *H. vulgare* individuals 95.8% of the characters were constant and 37,344 characters were parsimony informative. For the dataset consisting of 229 diploid *H. bulbosum* plus five *H. vulgare* individuals 96.3% of the alignment positions were constant and 34,383 characters parsimony informative. The alignment including only diploid *H. bulbosum* individuals had 97.5% invariable positions and 20,529 parsimony informative sites, while with only tetraploid individuals included it consisted of 98.8% constant alignment positions and had 8,821 parsimony informative characters. Adding the outgroups to the latter resulted in a dataset with 97.2% constant and 26,269 parsimony informative characters.

Our initial MP analysis including di- and tetraploid individuals and defining *H. vulgare* as outgroup resulted in 48 most parsimonious trees with a length score of 234,446 and a consistency index (CI) of 0.23 and retention index (RI) of 0.61. A schematic representation of the strict consensus of this tree is provided in Figure 2 and Figure S1 (all figures indicated with “S” are available online as supplementary materials). In this tree Libyan diploids are sister to all other *H. bulbosum* populations. The initial consecutively branching clades consist all of tetraploid individuals with Greek tetraploids being sistergroup to all diploid materials. The first clade in the latter group consists of Greek diploids harboring also a single tetraploid individual from the Greek Peloponnese area (FB19-001-1). The next branches consist of Albanian, southern Croatian and Greek populations which together are sistergroup to populations from Italy (including the islands of Sicily and Sardinia) and Spain.

**Figure 2.**
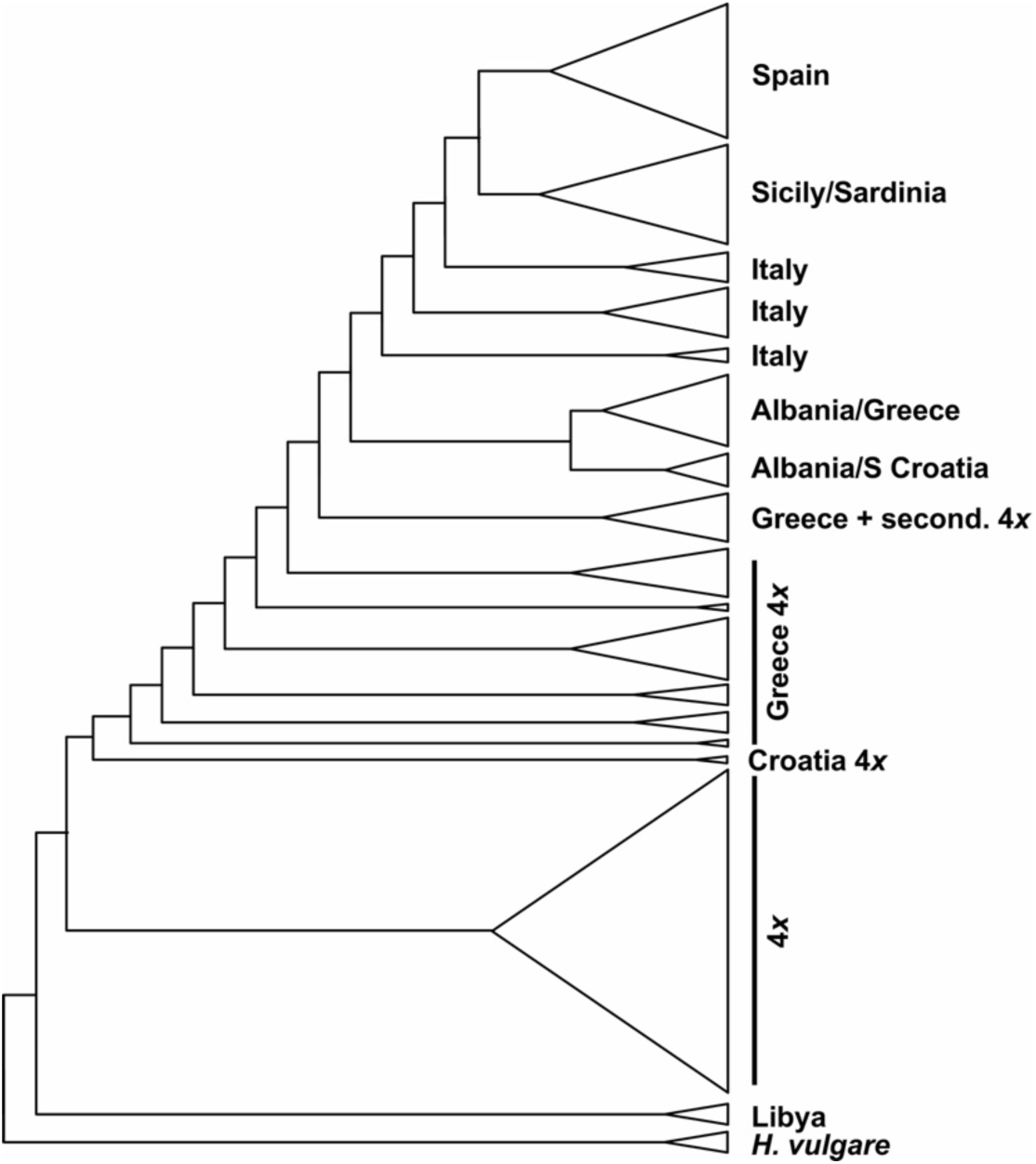
Scheme of the strict consensus tree topology derived from an MP analysis of GBS data including diploid and tetraploid *H. bulbosum* individuals. Barley (*H. vulgare*) was defined as outgroup. Tetraploids are depicted by “4*x*” and partly also their occurrence areas are given. For the diploid *H. bulbosum* populations the areas of origin are provided to the right.

In our second MP analysis including only diploid *H. bulbosum* populations together with the outgroup *H. vulgare* we obtained a single most parsimonious trees with a length score of 200,304 (CI = 0.24, RI = 0.56). A schematic representation of the tree together with the result of a neighbor-joining analysis is provided in Figure 3. Libyan individuals form the sistergroup to all other *H. bulbosum* populations and are genetically clearly different from them. The next branching clade consists of two individuals from Tarento in the southern Italian Apulia area. The rest of the tree consists of two clades, one harboring individuals from Apulia, and nested within them the populations from Albania, southernmost Croatia and Greece. The second clade consists of Italian populations from the mainland, partly intermingled with Apulian individuals and also the single population from Tunisia. Nested within these mainland Italy clades are the populations from the islands of Sicily/Sardinia, which together are sister to all Spanish populations. The Sicilian populations are nested within the ones from close-by Calabria, and the Sardinian populations within the Sicilian.

**Figure 3.**
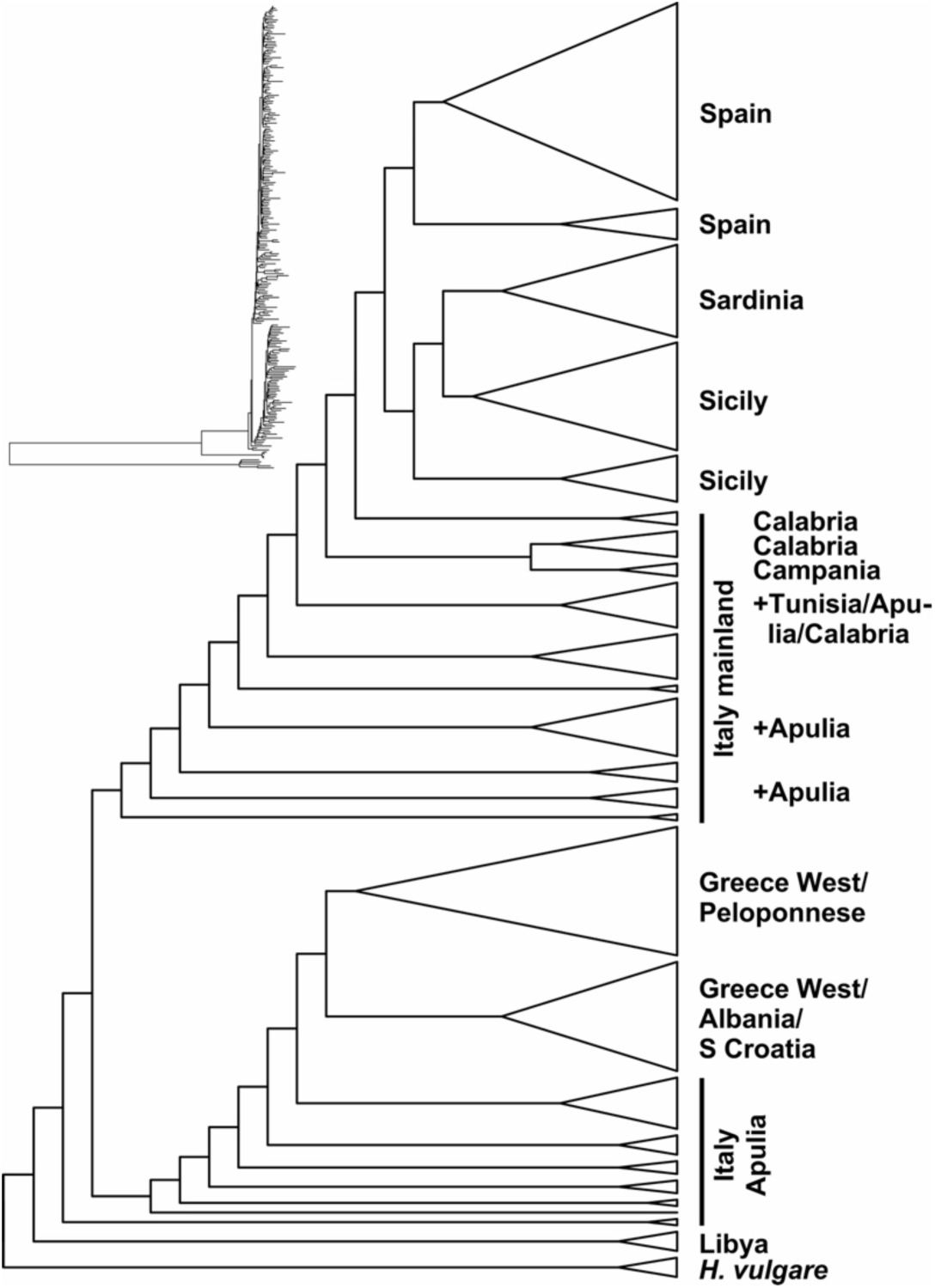
Scheme of the tree topology of the single most parsimonious tree derived from an MP analysis of GBS data of diploid *Hordeum* individuals. Barley was defined as outgroup. For the *H. bulbosum* populations the areas of origin are provided to the right. The insert on the upper left side provides branch lengths for this dataset derived from a neighbor-joining analysis.

When checking the robustness of our results by using IQ-TREE to calculate an ML-based phylogeny for the diploid individuals including *H. vulgare* as outgroup, we obtained a very similar tree topology as in the MP analysis of this dataset (Fig. S2). Excluding *H. vulgare* and instead defining the Libyan accession as outgroup resulted in MP (Fig. S3) and ML (not shown) analyses in the same tree topologies for the diploids as before.

In SVDQuartets analyses we partitioned the diploid individuals according to the areas they belong to or their (meta)populations. In both analyses (Fig. S4) the Libyan population resulted as sister to all other groups. When using areas as units we obtained a tree (Fig. S4A) where an Apulian subgroup is sister to Albanian/Croatian and Greek clades. They together are sistergroup of all other area groups, starting with Tunisia, which is sister to an Italian grade that harbors a Sicilian/Sardinian and Spanish clade. Using a finer subdivision of the Italian materials by defining population partitions (Fig. S4B) resulted essentially in the same overall topology with the main difference that in this case Sardinian accessions became sister to the Spanish clade. Support values are very high in the first analysis, while in the second analysis they are mostly high except for the branch uniting Calabria with the Italian islands and Spain (62%) and the sistergroup relationship of Sardinia and Spain (59%).

Regarding the tetraploids, MP analyses without an outgroup (Fig. S5) and with changing diploid taxa defined as outgroups were conducted. In all cases we excluded a secondary evolved tetraploid population from the eastern Peloponnese in Greece (FB19-001) and restricted the analyses to the main clade of tetraploids. Figure 4 summarizes the different trees obtained from these analyses. Generally, we found that geographically closer populations group (mostly) together. Thus, Tajikistan and Uzbekistan, Armenia and Turkey form stable clades that group mostly together with Israeli and Ukrainian plants. The single population from Morocco falls in between French and the Israeli/Ukrainian accessions. Populations from Greece, including the northern Aegean islands, group close to southern Croatian tetraploid populations, Bulgarian individuals, southern Aegean islands, and the populations from Istria in northern Croatia. Depending on the included outgroups, the tree topology flips or changes. In the case of *H. vulgare* as outgroup, Moroccan materials are sister to all other included tetraploids, which form an eastern (Ukraine to Tajikistan) and western clade (Bulgaria, France, and Croatia to Greece). Using the Libyan population instead of barley as outgroup results in southern Croatian populations being on the one hand sister to Greek materials and on the other to all other populations (Fig. 4C). Using Greek diploids as outgroup resulted in Greek tetraploids forming the first clade to branch off, followed by a grade consisting of southern Croatia, southern Aegean islands, Bulgaria, Istria, France, Morocco, and the western Asian populations. In cases where the included outgroups resulted in deviations from the topology of the unrooted tree without outgroup, we used these changes as indications for either introgression (Fig. 4D) or presence of old plesiomorphic characters retained in some regions of the tetraploids’ distribution areas (Fig. 4B, C).

**Figure 4.**
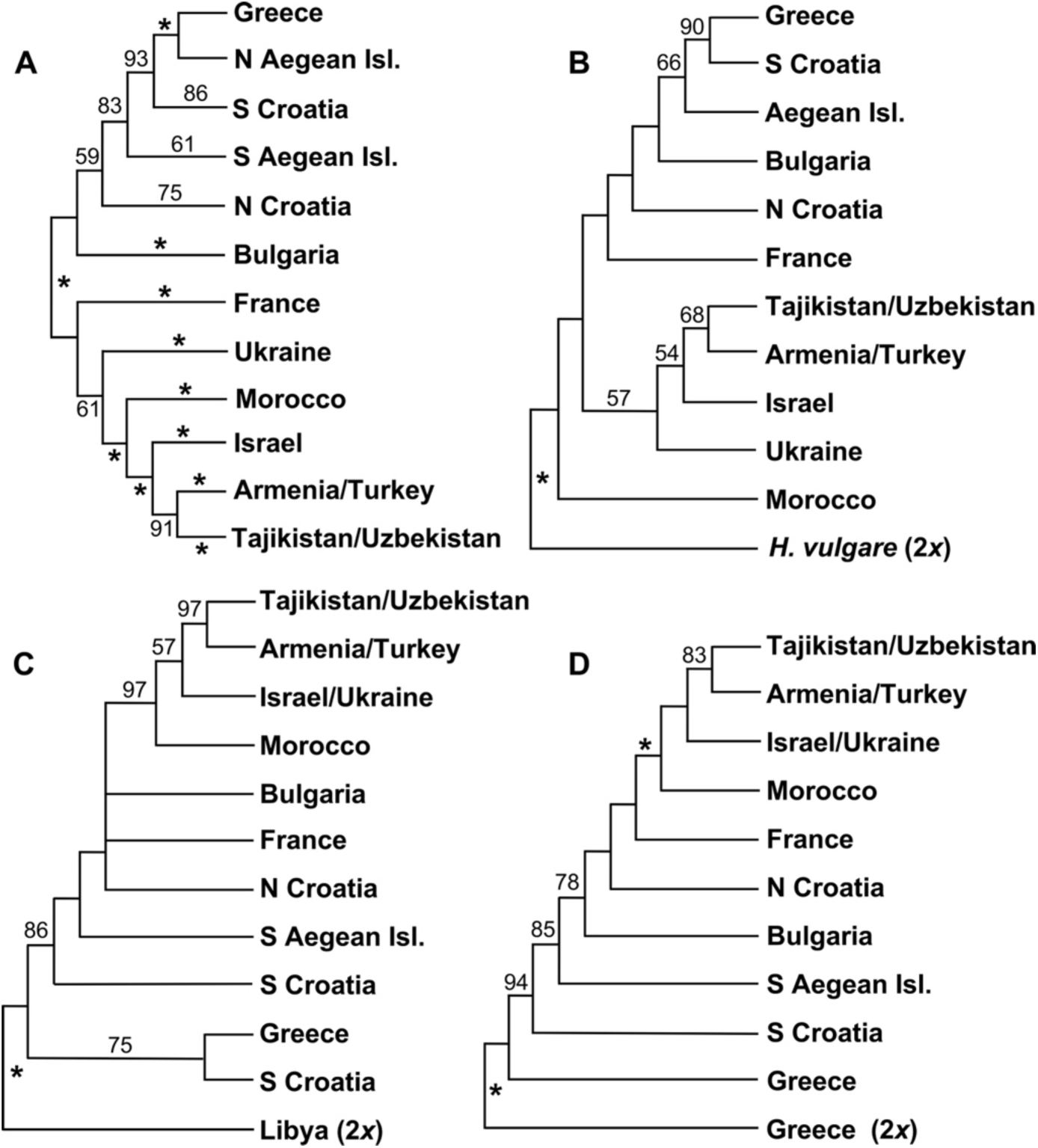
Schematic representation of the strict consensus trees derived from MP analyses of tetraploid *H. bulbosum* individuals. For the populations the areas of origin are given. **A** Unrooted tree including tetraploids only. **B** Tree rooted with diploid *H. vulgare* as outgroup. **C** Tree rooted with Libyan diploids as outgroup. **D** Tree rooted with Greek diploids as outgroup. Bootstrap support values are provided along the branches with asterisks indicating ≥99% support. In **A**, support values are given for the populations from specific areas, while in the other trees support values are provided only for the backbones of the trees.

### Population assignment analyses

For the diploid *H. bulbosum* dataset (consisting of 230 individuals and 5,121 SNPs), Bayesian assignment analyses by LEA and STRUCTURE identified K = 5 as the best representation of ancestral populations (Fig. 5). Libyan samples were assigned to a distinct ancestral group (brown in Fig. 5). The Greek populations form also a rather uniform group (green) with two Greek individuals (Peloponnese, Mani) carrying a proportion of about 9% (0.095 and 0.085) of the Libyan alleles. Apulian populations are largely assigned to the same group as the individuals from Greece but share in addition a fraction of alleles with other parts of Italy. The populations from the Italian islands of Sardinia and Sicily are falling in one group (mostly blue) with Sicilian individuals forming the connection to Spain. The Spanish cluster (mostly red) consists of individuals, which share similarities with Sicily. PCA of diploids (not shown) separated the Libyan population from the other individuals along PC2 and resulted in similar groupings as in the STRUCTURE plot.

**Figure 5.**
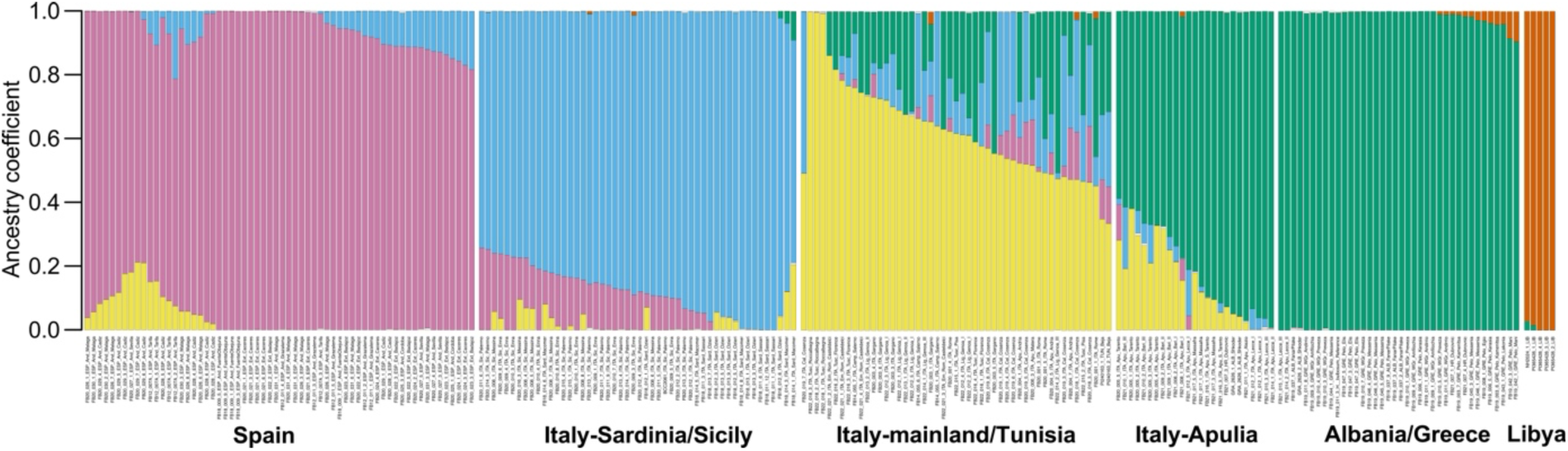
Plot of a Bayesian population assignment analysis conducted in STRUCTURE for diploid *H. bulbosum* individuals at K = 5.

For the dataset including di- and tetraploid *H. bulbosum* individuals, PCA clearly separated diploids from tetraploids and LEA and STRUCTURE indicated K = 11 as best representation for the populations. In Figure 6, we display the PCA results, with individuals represented as pie charts providing a color code for the fractions of their group assignment in STRUCTURE. Libyan diploids cluster within the tetraploids, while the secondarily evolved tetraploid individuals from the eastern Peloponnese cluster closely with their diploid progenitors (mostly yellow), but also exhibit signatures of introgression from the tetraploids from the same area. In the ancestral population bar plot (Figs. 6B, S6) these secondary tetraploids share alleles with diploids from western Greece (yellow, and additionally blue for individual FB19-001-1) and tetraploids from the Peloponnese area (light blue, green).

**Figure 6.**
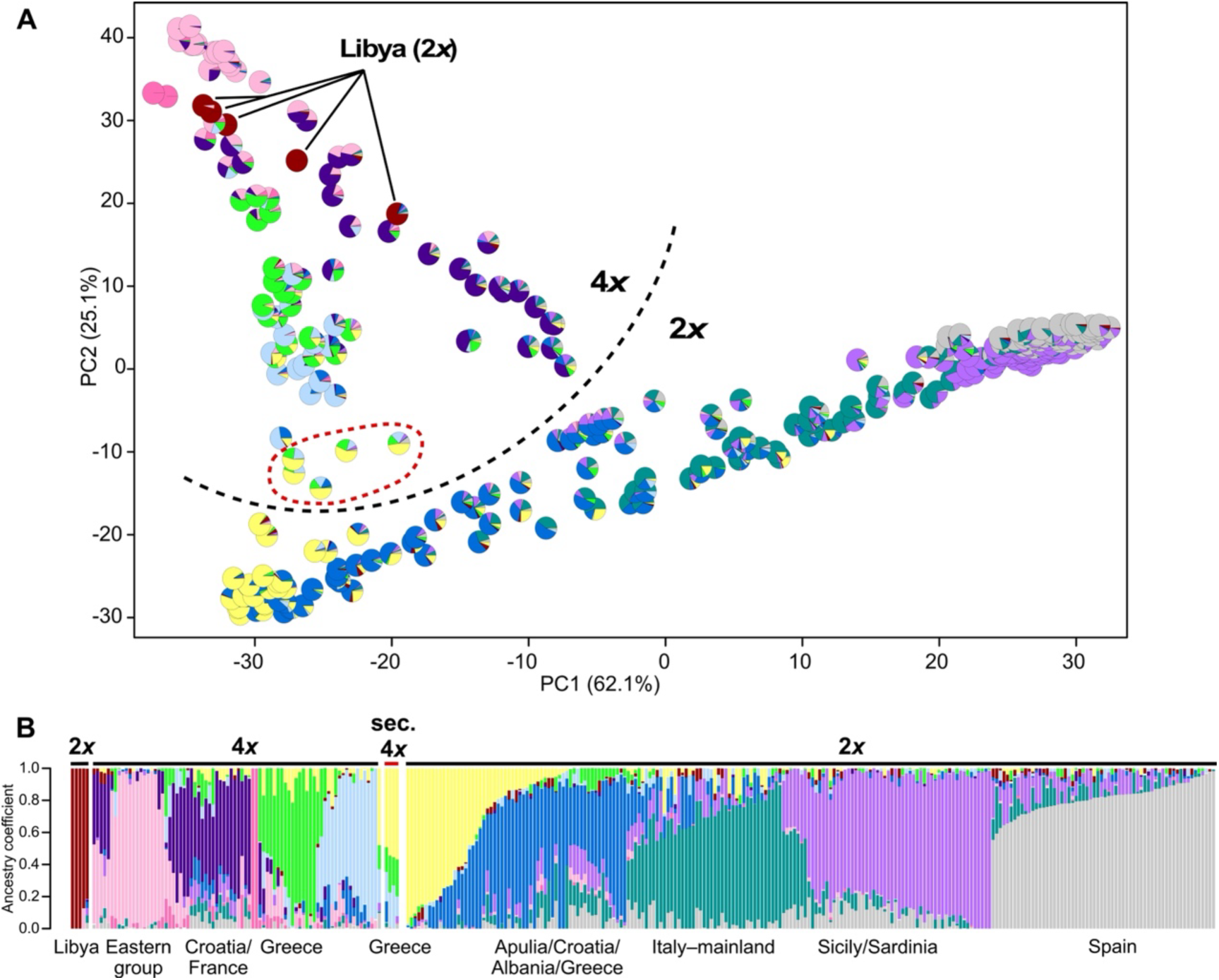
Results of the analyses of di- and tetraploid *H. bulbosum* individuals from (**A**) Principal Component Analysis (PCA) and (**B**) Bayesian population assignment analysis conducted in LEA at K = 11. Pie charts in the PCA plot reflect the proportions of ancestral population contributions according to the LEA analysis. Libyan diploids group in PCA within the tetraploid populations. The secondary evolved tetraploids from eastern Peloponnese (red dashed line) group close to their diploid progenitors (mainly yellow) in PCA and show in both analyses varying amounts of introgression by Greece tetraploids.

### Ecoclimatic niche modeling

Ecoclimatic niche modeling showed large areas of overlap of suitable habitat of the two *H. bulbosum* cytotypes, which comprised almost the entire Mediterranean Basin (Fig. 7). Thus, models predicted the possible common presence of both cytotypes in, e.g., Italy, Sicily, Sardinia, Tunisia, and along the southern parts of the Black Sea, indicating very similar ecoclimatic niche requirements. However, in other cases, the modelling outcome correctly reflected the mutual exclusive presence of cytotypes, as could be observed for Central Asia (only tetraploids present) and Spain (only diploids present). The clear geographic separation between the cytotypes in Greece, where diploids occur only west and tetraploids only east of the Pindos/Taygetos mountain ranges, can only be observed in the model of the diploids. Statistics of niche overlap were D = 0.574 and I = 0.852, respectively, i.e. ecoclimatic niches of diploids and tetraploids are rather similar.

**Figure 7.**
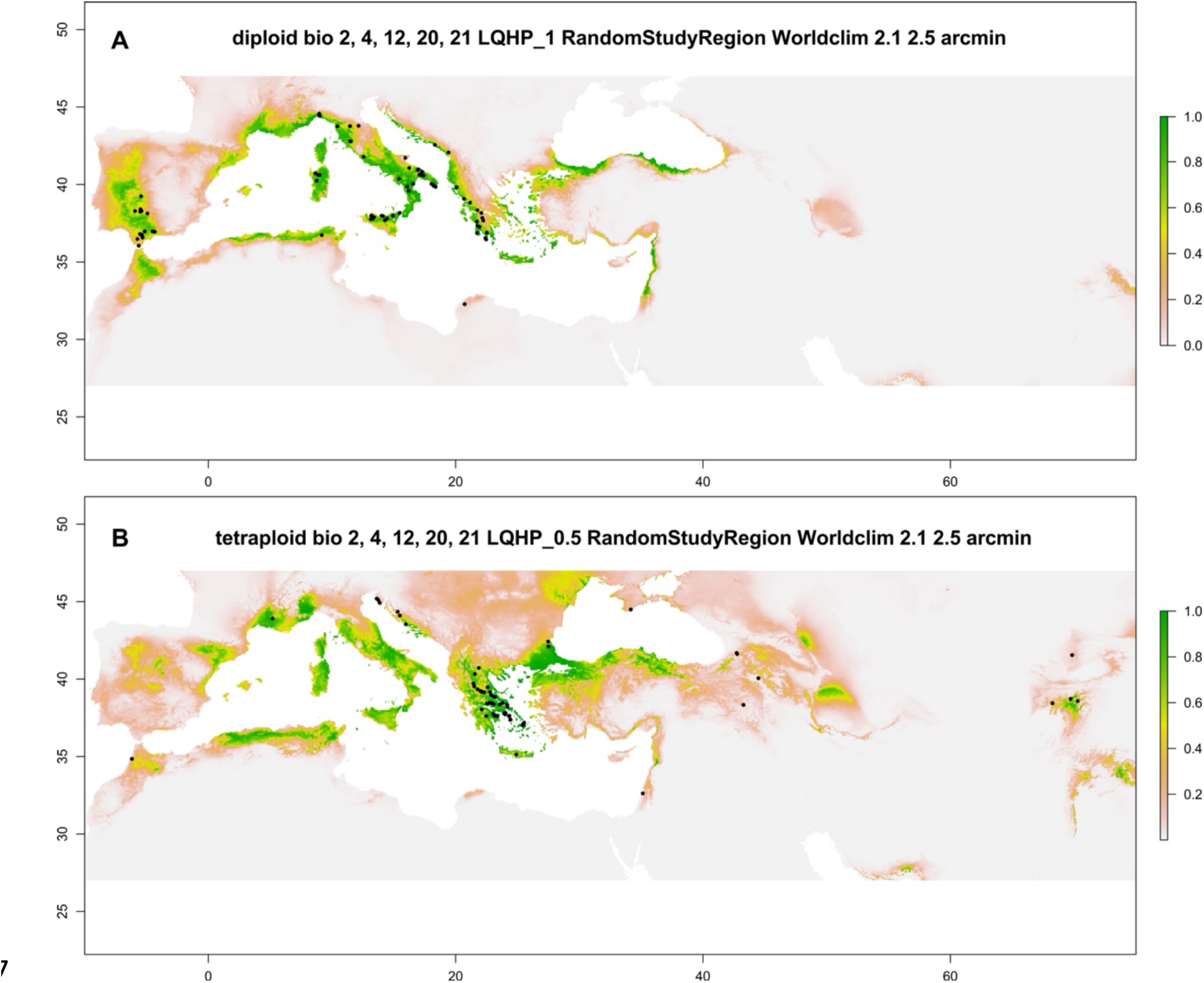
Ecoclimatic niche models for *H. bulbosum* cytotypes. The scale to the right reports the color code for habitat suitability, black dots within the maps depict the population collection sites. (**A**) Modeling of habitat suitability for the diploid cytotype based on the collection data of the analyzed populations. (**B**) Modeling of habitat suitability for the tetraploid cytotype based on the occurrence data for the analyzed populations.

## Discussion

### Population differentiation within diploid H. bulbosum

In many cases, the support values and consistency indices of our parsimony trees are relatively low within *H. bulbosum*. This is often the case in intraspecific analyses, where clades are usually young and genetic differences are not readily apparent. In addition, in outbreeding species such as *H. bulbosum*, gene flow and genomic recombination contribute to homoplasy in the datasets. Thus, low support values are to be expected and do not indicate shortcomings of the GBS method. Nevertheless, many of the individuals from single populations or from specific areas consistently group together and receive bootstrap support values >50%, and in ML analysis the backbone of the phylogeny as well as geographic groups received rather high support (Fig. S2).

The initial split in all phylogenetic trees separates Libyan individuals from all other *H. bulbosum* populations examined (Figs. 2, 3). This is also obvious in the PCA and STRUCTURE results (Figs. 5, 6). As the Libyan population is genetically very different from European *H. bulbosum*, we consider it a potentially important genotype within the species that is currently understudied. Occurrences of the species are reported from northern Cyrenaica, i.e. northeastern Libya (El Rabiai *et al*., 2010; Sherif & Mahklouf, 2020). Like other diploids of *H. bulbosum*, populations in Libya are patchily distributed and the species appears to be rare. The low representation of Libyan *H. bulbosum* in genebanks, in addition to its rarity, can be explained by the relative isolation of the country during the last half century, which has hindered systematic collections.

Within the mainly European group of diploid populations we found three major clades (Figs. 3, S2). Two individuals from the Apulian area of Tarento are sister to all other populations, which are then divided into a clade with Apulian, Albanian, Croatian and Greek populations, and a clade consisting of Italian (including Sicily, Sardinia and Tunisia) and Spanish populations. On the Balkan Peninsula, populations from Albania and southern Croatia are grouped together with western Greece. In mainland Italy, no clear geographic structure was found north of Puglia and Calabria, suggesting ongoing gene flow among the populations. The populations from the islands of Sicily/Sardinia are nested within the Calabrian populations as well as the Spanish individuals. In the latter group, a pattern similar to that in mainland Italy was found, with no clear separation of populations and areas (Fig. S2), suggesting high levels of gene flow between populations. This hypothesis is supported by the STRUCTURE and PCA analyses. STRUCTURE sorted individuals into five groups with the highest amount of diversity or ancient allele types found in mainland Italy (Fig. 5). The Spanish populations are more uniform in this analysis and, as in mainland Italy, the different allele combinations are not geographically sorted. In contrast, island populations from Sicily and Sardinia form one group, but are internally assigned to different allele types. Southeastern Italian populations from Apulia share large portions of their alleles with *H. bulbosum* individuals east of the Adriatic Sea. The same pattern occurs in Spain, where local populations share ancestry with Sicilian individuals, but with different proportions of ancient units (Fig. 5). We interpret these patterns as indicating ancestor-derivative relationships between populations from Apulia and Albania/Greece on the one hand, and Sicily and Spain on the other. Similarly, mainland Italy shares its pattern with the Tunisian population, which we also interpret as close relationship.

### Phylogeography of diploid H. bulbosum

The split between the lineages leading to *H. bulbosum* and *H. vulgare* has been dated to 4.5 Mya (Brassac and Blattner, 2015). Since the populations of wild *H. vulgare* occur in the Fertile Crescent and eastward and the basal lineages of the diploid *H. bulbosum* occur in Libya and southern Italy, we hypothesize that both species began to separate in the eastern Mediterranean after the end of the Messinian salinity crisis (5.3 Mya). Subsequently, the eastern population evolved into barley, while bulbous barley evolved in the west.

Our phylogenetic analyses consistently showed that Libyan individuals of *H. bulbosum* are sister to all other populations of the species and are genetically quite distinct (Figs. 3, 5, 6). A genome-based estimate for the split between these *H. bulbosum* groups arrived at about 2 Mya and at 1.5 Mya for the origin of tetraploids (Feng *et al*., 2025). Thus, the distribution of *H. bulbosum* on the northern and southern shores of the Mediterranean cannot be attributed to migration across the Mediterranean basin during the Messinian event, but must have involved dispersal across the Mediterranean or migration through the Levant.

In the European populations of the species, Apulia, the southeastern region of Italy, was the starting point (Fig. S7) for the colonization of the Balkan Peninsula, as Albanian and Greek *H. bulbosum* populations are nested within individuals from Apulia. Within Greece, we see a stepwise southward migration from western Greece through the western parts of the Peloponnese (Figs. S2). The pattern is less clear for mainland Italy, but even here Apulian individuals are interspersed among populations occurring further north (Figs. 3, S2), which indicates an ‘out of Apulia’ scenario (Fig. S7). Otherwise, most Italian clades are of mixed origin, due to gene flow between populations in the central and northern parts of mainland Italy. Southwestern Italy was the starting point for the colonization of Sicily and Spain, as Sicilian and Spanish populations are nested within the Calabrian genotypes. Similarly, Sardinian populations are nested within those from Sicily (Figs. 2, S2), indicating that Sardinia was reached from Sicily. For Tunisia, we postulate long-distance dispersal (LDD), as Tunisian individuals are nested within populations from mainland Italy. While LDD is frequent in *Hordeum* (Blattner 2006), in this case, the SVDQuartet analyses might suggest a different interpretation, as they placed the Tunisian individuals as sister to the majority of the Italian populations (Fig. S4), which, taking into account also the Libyan population, could be interpreted as an origin of *H. bulbosum* in North Africa followed by dispersal to Europe. However, to test this hypothesis, additional populations from North Africa would have to be included. While the Spanish population is monophyletic, there is no clear geographic pattern within Spain (Figs. 6, S2).

### Multiple origins of tetraploids and gene flow among cytotypes

Di- and tetraploid cytotypes of *H. bulbosum* are morphologically quite variable and not clearly distinguishable without genome size determination or chromosome counting (Jørgensen, 1982). The main difference we have observed in the field is mostly related to population size. While diploids, even if quite common in an area, mostly occur in smaller, patchily distributed stands, tetraploid populations tend to be larger and sometimes form almost continuous stands over tens of kilometers (e.g., in Istria or central Greece). However, there are exceptions, such as diploids occurring in large numbers in the valleys of northern Sicily and the Pollino Mts. of Italy, and tetraploids growing in widely dispersed small populations on some of the Aegean islands.

Initial phylogenetic analyses including diploid and tetraploid individuals always resulted in an individual from a population from the eastern Peloponnese in mainland Greece (FB19-001-1) being grouped separately from all other tetraploids within diploid individuals from the western Peloponnese (Fig. S1). PCA and population assignment results also reflect this relationship (Figs. 5, 6). This strongly suggests that this autotetraploid population from the eastern Peloponnese evolved independent of all other tetraploids from local Greek diploids. Comparative sequencing of the genomes of the two tetraploid types from Greece as part of an *H. bulbosum* pan-genome project confirmed this finding, as the genomes are clearly distinct and the secondary tetraploid is very young and shares large parts of its genome with Greek diploids (Feng *et al*., 2024). Other individuals from the same population (FB19-001-2 to 5) were found at an intermediate position in the phylogenetic trees. Population assignment analysis suggests that they share alleles with tetraploids from the Peloponnese, a result supported by their position in the PCA plot (Fig. 6). We interpret this as an indication of ongoing gene flow between these separately evolved autotetraploids and members of the main clade of tetraploids. Over time, this leads to assimilation of the newly formed tetraploids into the pre-existing older gene pool (Soltis & Soltis, 1992, 1999). We excluded these secondary tetraploids from subsequent phylogenetic analyses of the other tetraploids to avoid the influence of their mixed heritage on the topology of the phylogenetic trees.

The initial phylogenetic analysis, including diploids and tetraploids, showed that all diploid populations, except the one from Libya, grouped within a grade formed by tetraploids (Fig. 2). This would imply that diploids re-evolved from the tetraploid cytotype, with diploidization resulting in half the chromosome number and genome size of the tetraploids. This scenario seems highly unlikely, especially since we found that the phylogenetic trees of the tetraploids changed when different diploids were included as outgroups (Fig. 4). In parsimony analyses, this behavior is typical when hybrids or genomically introgressed individuals are involved. Since *H. bulbosum* is autotetraploid, we interpret the changing results as an indication of local gene flow between different di- and tetraploid individuals, which influences the genome composition of the involved tetraploids and thus the outcome of phylogenetic analyses. Therefore, we assume that the position of diploids within tetraploid populations in Figure 2 is an artifact caused by this effect and not by diploidization.

An analysis of tetraploids without a diploid outgroup (Fig. 4A) revealed several sistergroup relationships. Thus, Armenian and Turkish populations are grouped together, as well as Tajik and Uzbek, Israeli and Ukrainian, and Greek and southern Croatian populations. The Croatian tetraploids occur as a northern and southern clade or grade, formed by mainly Istrian individuals (north) versus the population from the Croatian Split area (south). Also within the populations from Greece two clades occur. One consists of the mainland populations plus the nearby northern Aegean islands (Evia, Kea), while the other consists of the southern Aegean populations from Crete and Naxos.

Inclusion of *H. vulgare* as an outgroup resulted in the Moroccan population being sister to two main clades, one consisting of the eastern populations from the Levant, northern Black Sea area and western Asia, and the other with the mainly western populations from Bulgaria, Croatia, France and Greece (Fig. 4B). Using the Libyan diploid *H. bulbosum* as an outgroup, two main clades were obtained (Fig. 4C) with individuals from southern Croatian populations forming the first branches in both. When Greek diploids were included as an outgroup, Greek tetraploids followed by southern Croatian populations formed the first successively diverging clades (Fig. 4D). In general, the internal relationships among the tetraploid populations, i.e. their relative position to each other, remain quite stable in all analyses. What changes with different outgroups are the basal positions in the trees, i.e. the branches that define the first sistergroup relationships. We interpret these inversions in the trees as indications of, first, local gene flow between diploids and tetraploids in certain areas, particularly in Greece, where the largest contact zone between both cytotypes exists (Fig. 1; Katznelson & Zohary, 1967). The detection of a secondary tetraploid in this area and its hybridization with the older stock of tetraploids supports this interpretation (see above). Second, in the case of the Moroccan and Croatian populations, we currently assume that they have retained a fraction of plesiomorphic “Libyan-like” characters from the time of their origin, which have not been swamped out by subsequent introgression. This hypothesis is supported by the result of the PCA and admixture analysis in LEA (Fig. 6), which show that the tetraploid populations group with Libyan diploids, while the Greek tetraploids in contrast are closer to their local diploid neighbors. Katznelson & Zohary (1967) hypothesized introgression of Near East tetraploids by another *Hordeum* species in this area. Based on distribution and possible hybridization abilities this could have only been *H. vulgare* or an extinct genotype from the vulgare/bulbosum lineage. In our data we find this eastern group of *H. bulbosum* populations strongly supported and grouping together with Morocco (Fig. 4). This could be an indication that these populations have a different genome composition (Fig. 6) in comparison to most other tetraploids. However, as the populations are also geographically close, we cannot decide if this is a result of earlier introgression or common ancestry.

Cauderon & Cauderon (1956) mentioned the presence of tetraploid *H. bulbosum* in France without further specification, while Ortiz *et al*. (1984) reported two mixed populations of di- and tetraploids in southern Spain, and Jørgensen (1982) found a triploid plant in the latter area. The single population from France that we sampled was indeed tetraploid, while in Andalusia we found only diploid individuals. We interpret the mixed presence of both cytotypes in Spain in the middle of otherwise only diploid populations as a strong indication for another secondary origin of the tetraploids and continuous admixture between both cytotypes resulting in triploid hybrids (Jørgensen, 1982). Tetraploids seem to be quite rare in Spain, but their occasional presence indicates that the formation of autotetraploid cytotypes is an ongoing process in *H. bulbosum*. The French tetraploids in our phylogenetic trees are always close to the tetraploid populations from northern Croatia, indicating that they belong to the main stock of tetraploids that reached the northwest of the species’ distribution area. They show no close relationship to the diploid cytotypes from Liguria in northern Italy, which are geographically their closest neighbors. In this case, we found no evidence of gene flow between neighboring cytotypes.

Tetraploids from Morocco always group close to or in between tetraploid populations from the Levant/Ukraine and France. They show no closer relationship to Spanish diploids, from which they are separated only by the Strait of Gibraltar, or to the Tunisian diploid population. It should, however, be emphasized that (i) we analyzed here a genebank accession and no population sample collected in the wild, (ii) that neither Katznelson & Zohary (1967) nor Jørgensen (1982) reported any tetraploids from Morocco, and (iii) di- and tetraploid *H. bulbosum* from North Africa are underrepresented in our dataset and generally very rare in genebank collections. Therefore, conclusions about relationships, ploidy-level distribution, and colonization routes of North African populations are premature at this time.

### Geographic separation of cytotypes

Jørgensen (1982) already reported a clear separation of di- and tetraploid cytotypes of *H. bulbosum* by the north-south extending mountain systems of central Greece. Our analyses show that the pattern of strong cytotype separation extends even further north. The diploids occur in the westernmost parts of Greece (Jørgensen, 1982) and reach as far north as Albania and the Dubrovnik area of southern Croatia. Further north, no diploids were found in Croatia and Slovenia, consistent with an earlier record of exclusively tetraploid cytotypes along the Adriatic coast of former Yugoslavia (Papeš & Bosiljevac, 1984). Essentially, the tetraploid populations of the Balkan Peninsula are distributed between northern Croatia/Slovenia, Serbia, and North Macedonia (Papeš & Bosiljevac, 1984), where they connect with the tetraploid populations of Greece on the one hand, and on the other, eastward to the Bulgarian and Black Sea stocks of the species. The tetraploid *H. bulbosum* population of southern France is an isolated northwestern outlier.

It is interesting to note that the independently evolved Greek tetraploid occurs in an area otherwise populated only by tetraploid cytotypes, rather than in the western parts of Greece where its diploid ancestral populations thrive. This, together with the ephemeral occurrence of tetraploids in Spain, suggests that newly evolved autotetraploids of *H. bulbosum* may not be able to establish stable populations when surrounded and outnumbered by the diploid cytotype, a phenomenon known as minority cytotype exclusion (Levin, 1975). Because both cytotypes largely share the same ecoclimatic niche (Fig. 7), they cannot diverge on a smaller geographic scale, which could otherwise lead to parapatric ecology-driven secession. In contrast, the geographic separation of the secondarily formed tetraploids from their parental population in Greece must have allowed them to escape the pollination pressure of diploids and the resulting mostly sterile triploid offspring. In Spain, where only diploids occur, the ephemeral occurrence of triploids (Jørgensen, 1982) is a strong indication for this mechanism. In the case of an independent secondary establishment of a polyploid population, as seen in Greece, interbreeding with other pre-existing tetraploids (Fig. 6) leads to an enrichment of genetic diversity within the tetraploid cytotype (Soltis and Soltis, 1999), providing an avenue for increasing the genetic diversity of polyploids in addition to mutations and genomic recombination.

## Conclusions

Our study provides a detailed overview of di- and tetraploid *H. bulbosum* populations, their connectivity and historical biogeography. Extending previous work, we were able to map the border zone between the two cytotypes on the Balkan Peninsula in more detail. We found that diploids occur only along the Adriatic/Ionian Sea between Dubrovnik (southern Croatia) in the north and the Mani peninsula of the Peloponnese (Greece) in the south. Further north along the Adriatic coast and east into western Asia, only tetraploids occur. We also confirmed the presence of tetraploid *H. bulbosum* in southern France. Otherwise, only diploid *H. bulbosum* is found in the central and western parts of the Mediterranean basin. The distribution of both cytotypes along the southern rim of the Mediterranean is currently unclear. Libya harbors a unique diploid genotype that differs from all other analyzed, the Tunisian accession is closely related to Italian populations, and a tetraploid from Morocco belongs to the main stock of tetraploid accessions. Thus, it is possible that both cytotypes co-occur in North Africa. However, this assumption is based on only a few accessions. Additional population sampling and genotyping seems necessary to draw conclusions for the species with respect to North Africa.

At the diploid level, we identified the southernmost Italian populations as the source for the colonization of the European part of the central and western Mediterranean. Thus, populations in southeastern Italy were the starting point for the colonization of the Balkan Peninsula. In Greece, we can trace the colonization route of *H. bulbosum* from Albania/northwestern Greece southward to the Peloponnese. Similarly, from Calabria (southwestern Italy) we inferred colonization through Sicily to Sardinia. Spain was also reached from Calabria. In contrast to Greece and southern Italy, we found no clear geographic structuring of genetic diversity in Spain and the central and northern parts of mainland Italy, indicating gene flow within Spanish and Italian populations and areas. Ecoclimatic niche overlap of both cytotypes is pronounced in the central parts of the Mediterranean basin, leading us to conclude that the exclusion of the minority cytotypes is the likely cause of the observed geographic distribution pattern. This implies that autotetraploids, after their first origin, essentially shared the same ecological niche with their diploid ancestors, without barriers to gene flow. The establishment of tetraploids was therefore only possible at the edge of the diploid range. Once a tetraploid population was successfully established, the new cytotype could not invade areas already populated by diploids, but had to expand eastward. Subsequently, the same mechanism prevented further eastward expansion of the diploids, resulting in the sharp geographic separation we see today. Autotetraploid formation is an ongoing process in *H. bulbosum*, as we found two independent origins of the tetraploid cytotype in our dataset, and short-lived tetraploid occurrences have been reported from Spain. The genome composition of the main stock of tetraploids shows similarities to the Libyan diploid genotype, indicating that they originated relatively early in the history of the species. Genomic introgression of tetraploids by local diploids was demonstrated for Greece, resulting in closer relationships of Greek tetraploids to local diploids than to diploids from other areas. Since autopolyploids readily form in the species, future breeding efforts can be made at the diploid level followed by genome doubling, which seems easier than dealing with the high allelic diversity present in the tetraploid cytotype.

## Acknowledgements

The authors thank Petra Oswald for support in the lab, Susanne König for GBS library preparation and sequencing, Danuta Schueler and Anne Fiebig for support with sequence data curation and submission. We acknowledge the support of the countries of origin for collection permits and thank Reinhard Fritsch and the IPK and USDA genebanks for the plant materials provided. Financial support through institutional funding by IPK is acknowledged.

## Data accessibility

The raw data of the GBS sequences used in our analyses can be accessed through the European Nucleotide Archive (ENA) project PRJEB73823.

## Author contributions

FRB designed the research. FRB, ABS, LS and AKA collected *H. bulbosum*. CK grew the plants and determined genome sizes. SP analyzed ecoclimatic niches. AH conducted sequencing. FRB and DH analyzed GBS data. FRB wrote the manuscript with input from all authors.

## Supporting information

The online version of this article provides the following supplementary information:

Supplementary file 1: **Figures S1 to S7 with extended phylogenetic trees, Bayesian population assignment plot, and colonization scenario for diploid *H. bulbosum***

Supplementary file 2: **Table S1 with details of the studied *H. bulbosum* and *H. vulgare* individuals**

## Supplemental figures online

**Figure S1.**
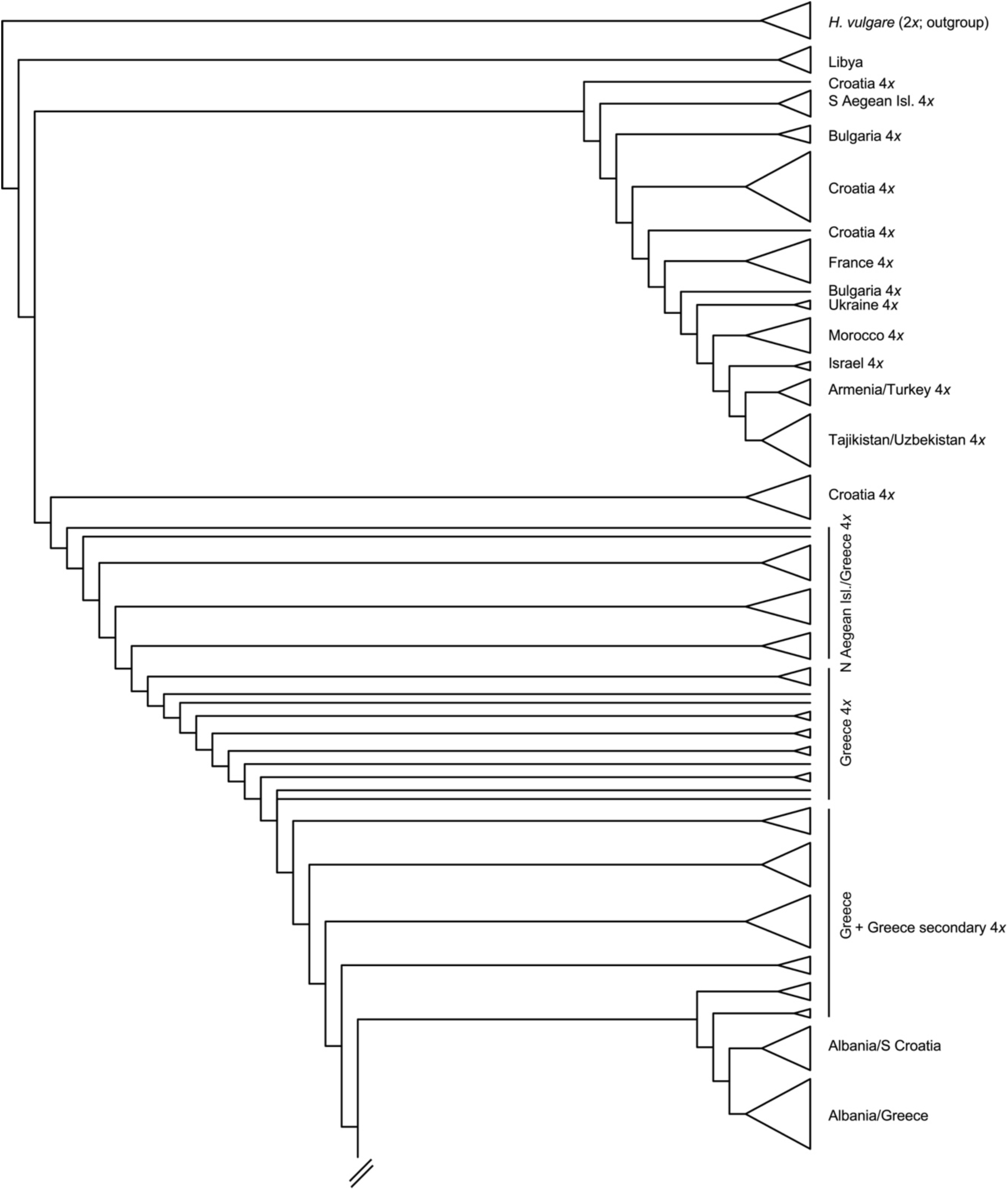
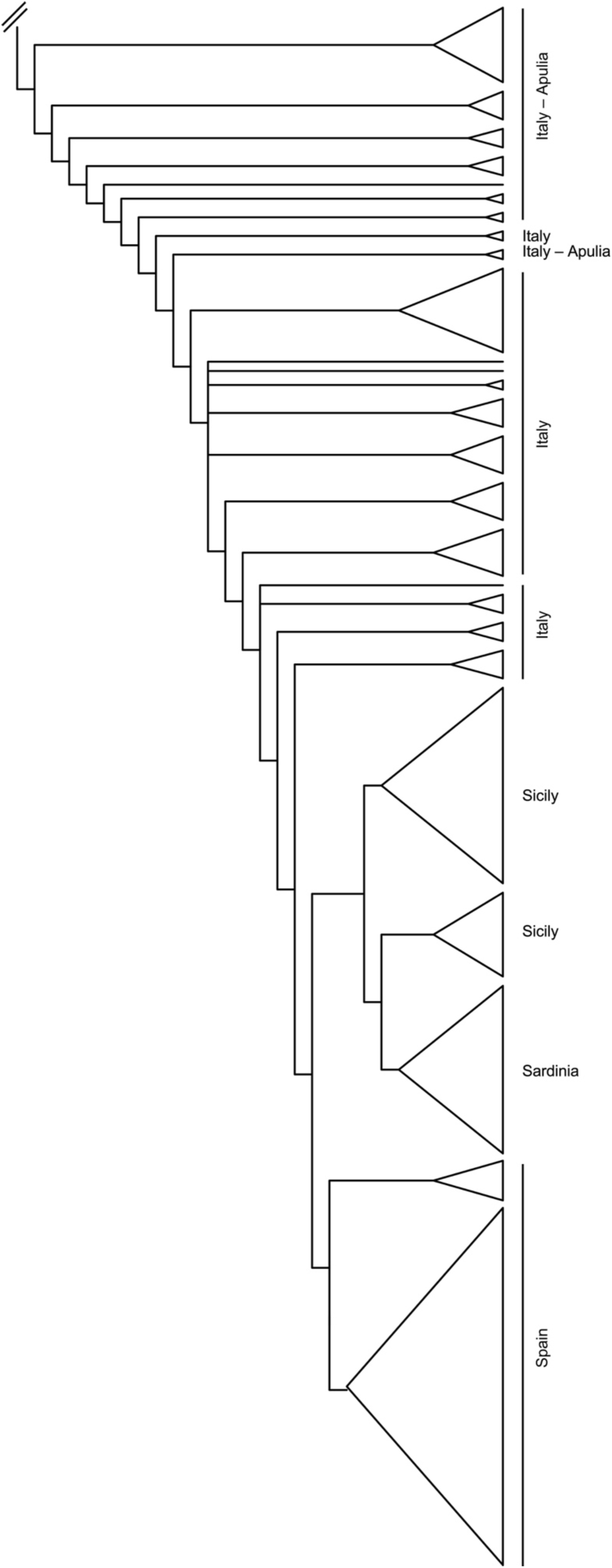
Scheme of the strict consensus tree of 48 most parsimonious trees resulting from an MP analysis of GBS data of diploid and tetraploid cytotypes of *H. bulbosum*. *Hordeum vulgare* was defined as outgroup. Tetraploid populations are indicated by “4*x*”.

**Figure S2.**
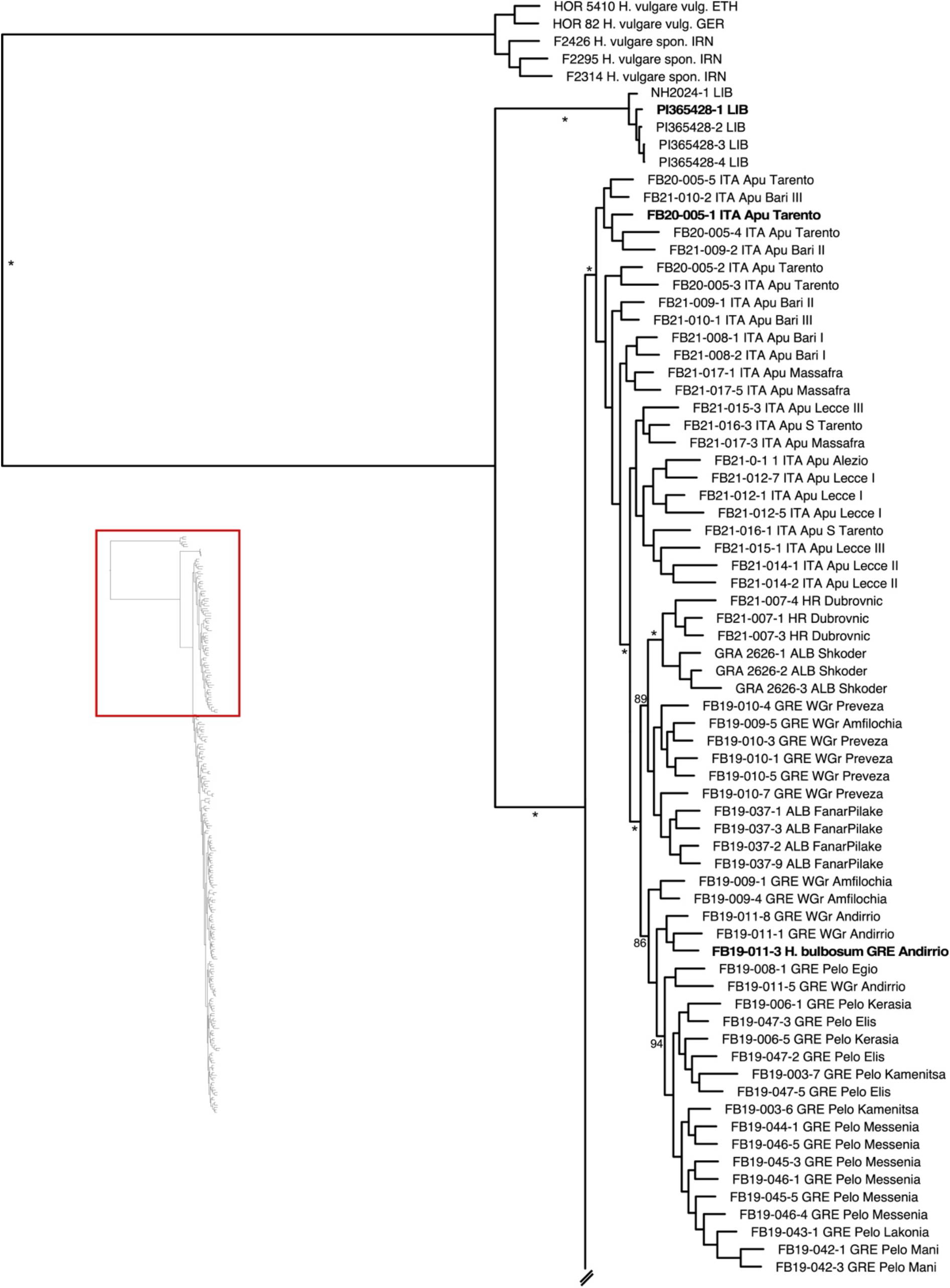
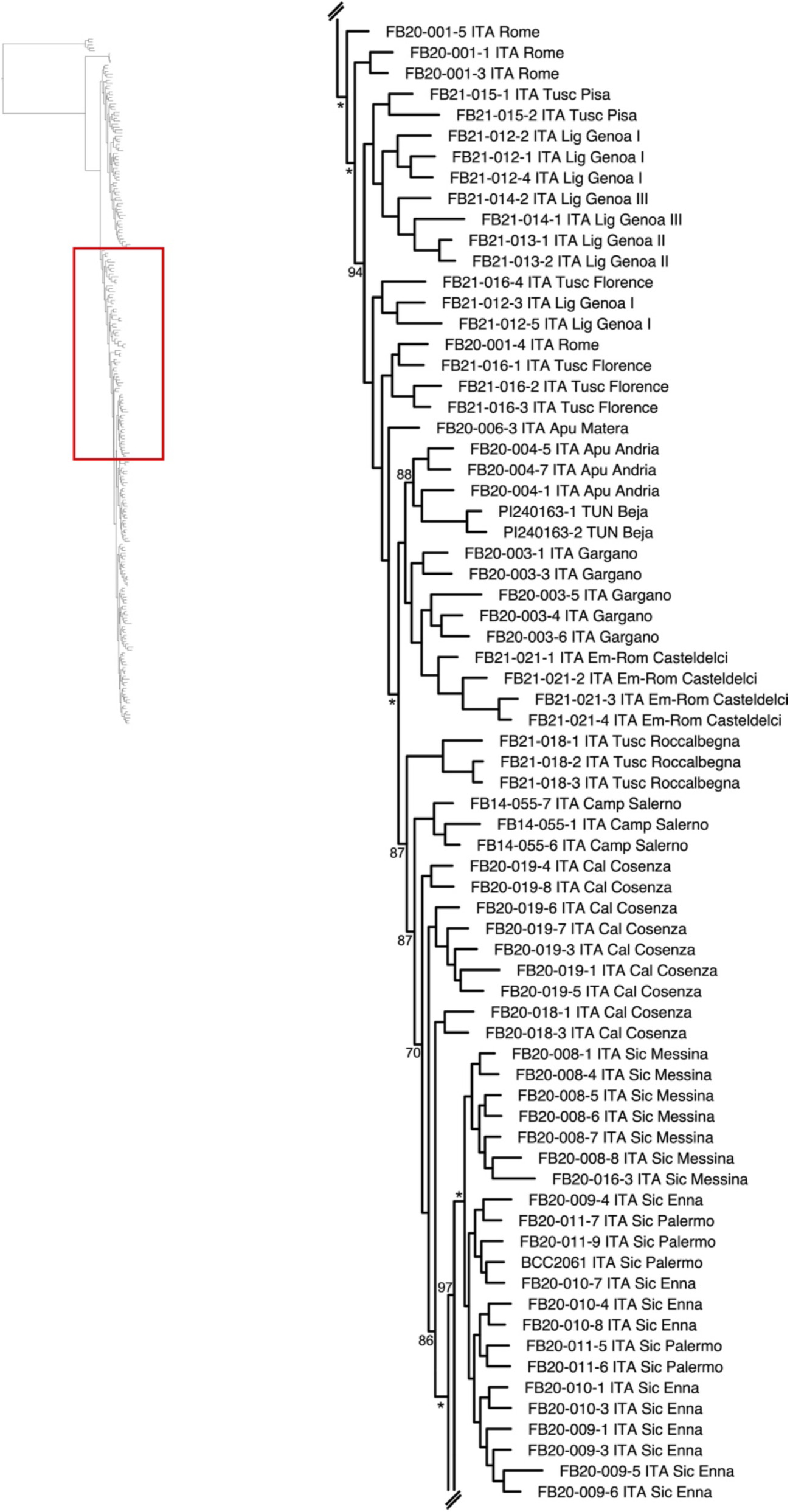
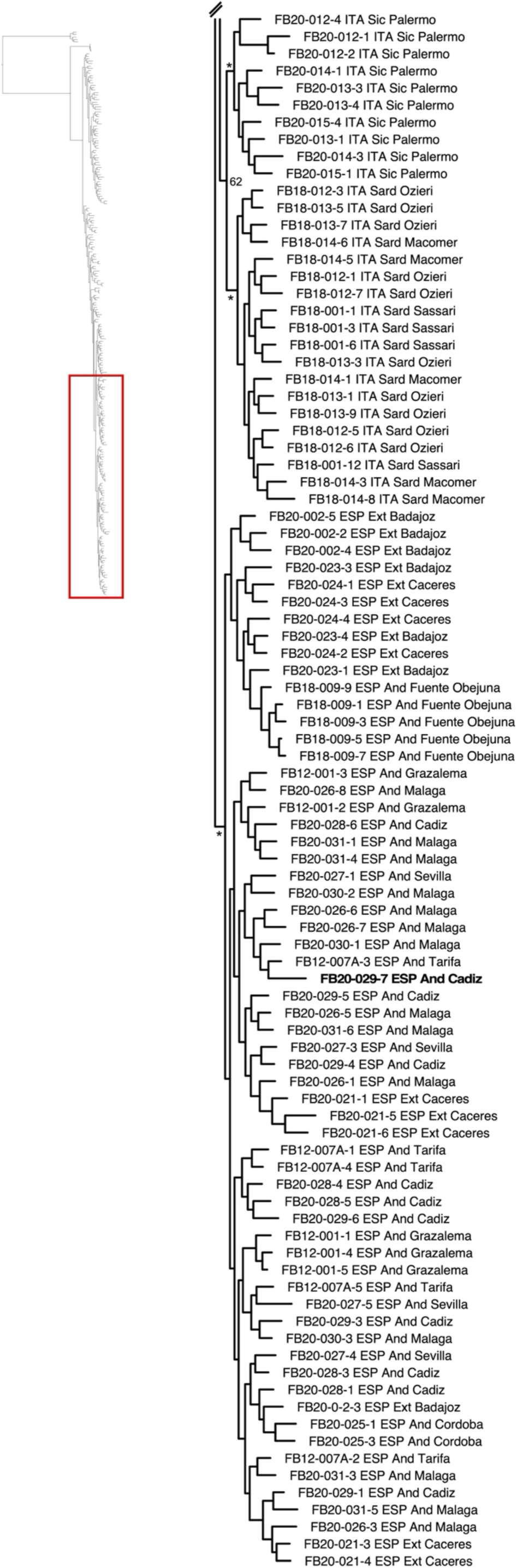
Phylogenetic tree resulting from an ML analysis of GBS data of diploid cytotypes of *H. bulbosum*, including *H. vulgare* as outgroup. Bootstrap values (%) for the tree backbone are given along the branches with asterisks indicating ≥99% support. Genomes of individuals provided in bold face were sequenced for a pan-genome of *H. bulbosum* (Feng *et al*., 2024).

**Figure S3.**
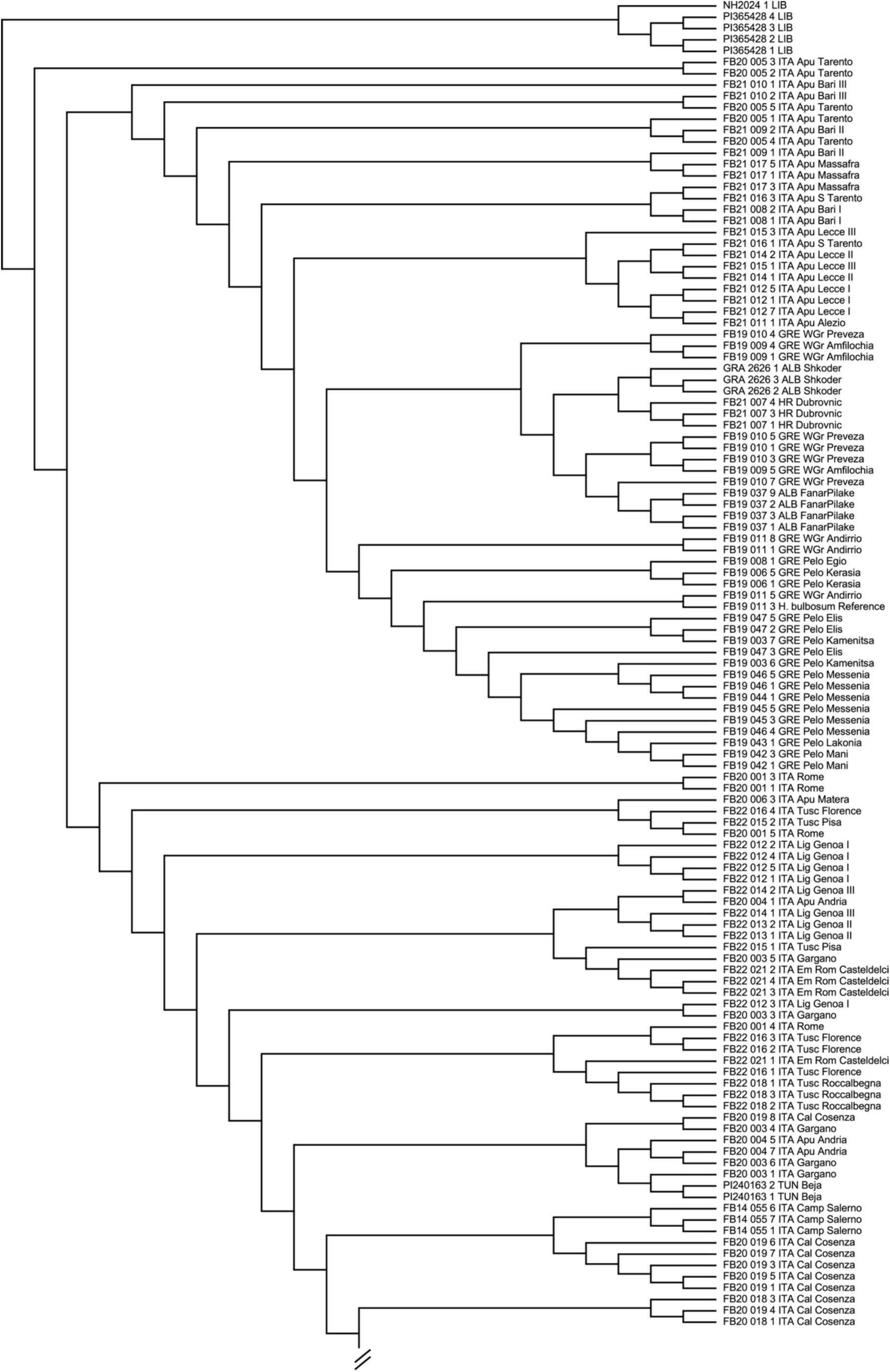
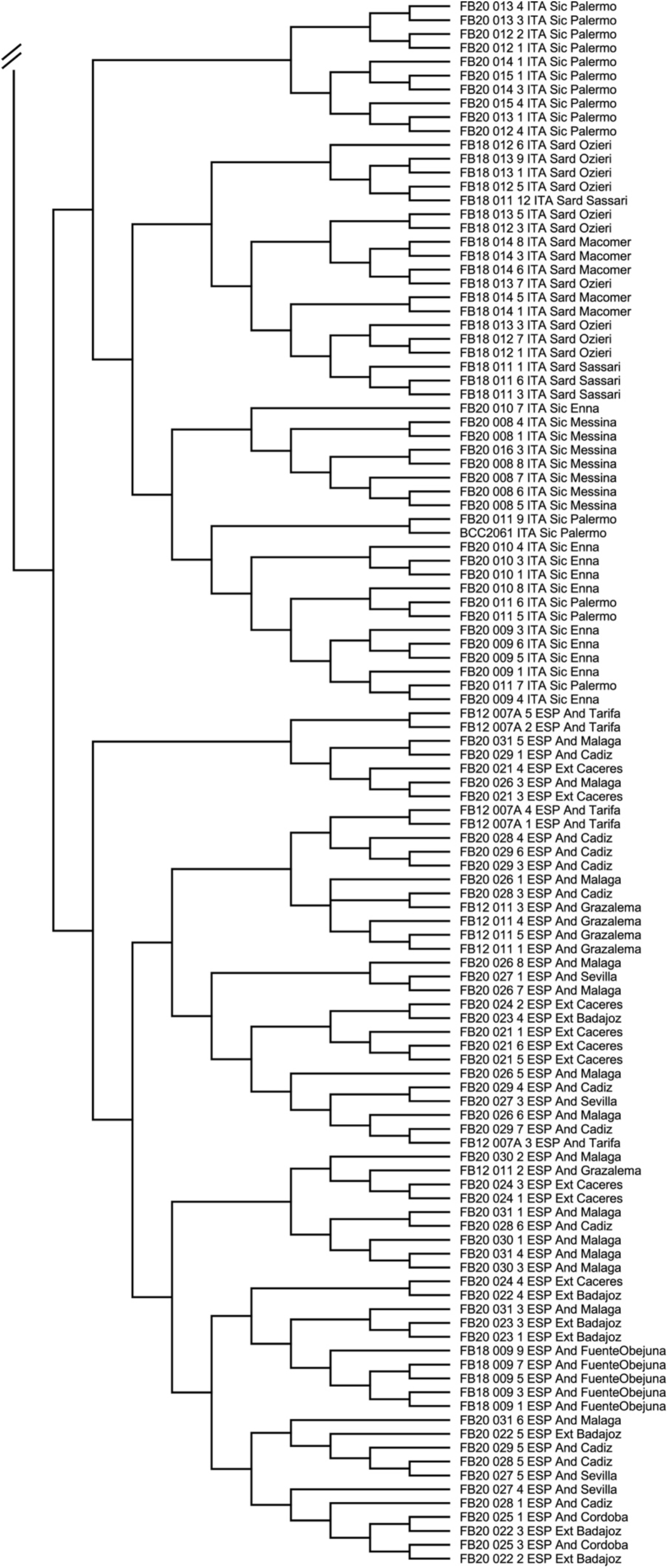
Strict consensus tree of two MP trees resulting from an MP analysis of GBS data of diploid *H. bulbosum* cytotypes. The Libyan individuals were defined as outgroup.

**Figure S4.**
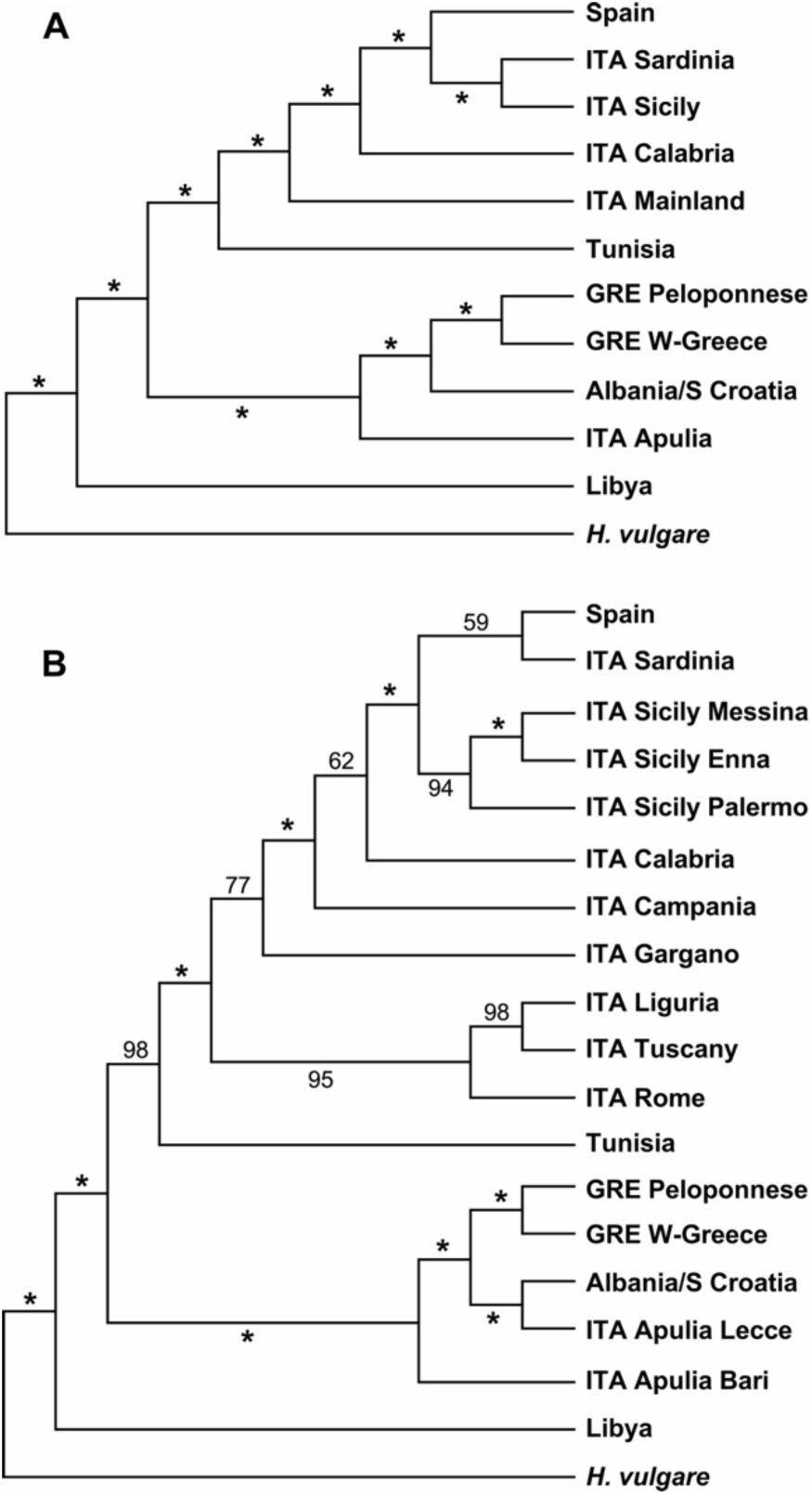
SVDQuartets-derived multispecies-coalescent trees based on the GBS dataset of diploid *Hordeum bulbosum*. *Hordeum vulgare* individuals were defined as outgroup. The dataset was partitioned according to areas (**A**) and populations (**B**). Bootstrap support values are given along the branches. Asterisks indicate values ≥99%.

**Figure S5.**
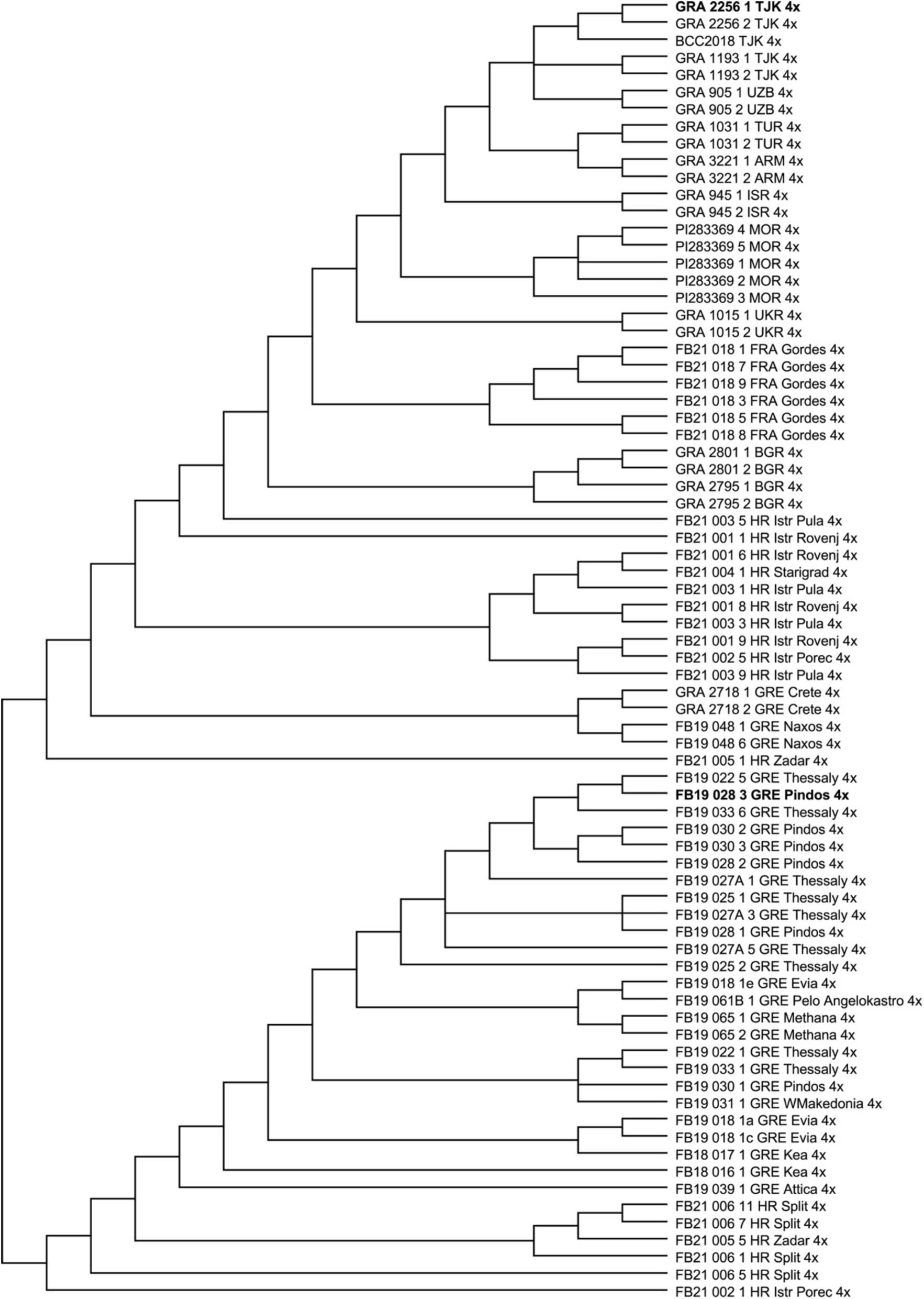
Unrooted strict consensus tree of 36 parsimony trees resulting from an MP analysis of GBS data of tetraploid individuals of *H. bulbosum*. No outgroup was included. Genomes of individuals provided in bold face were sequenced for a *H. bulbosum* pan-genome (Feng *et al*., 2024). The genome of the secondary evolved tetraploid FB19-001-1 was also sequenced, but is excluded from the phylogeneytic analysis here.

**Figure S6.**
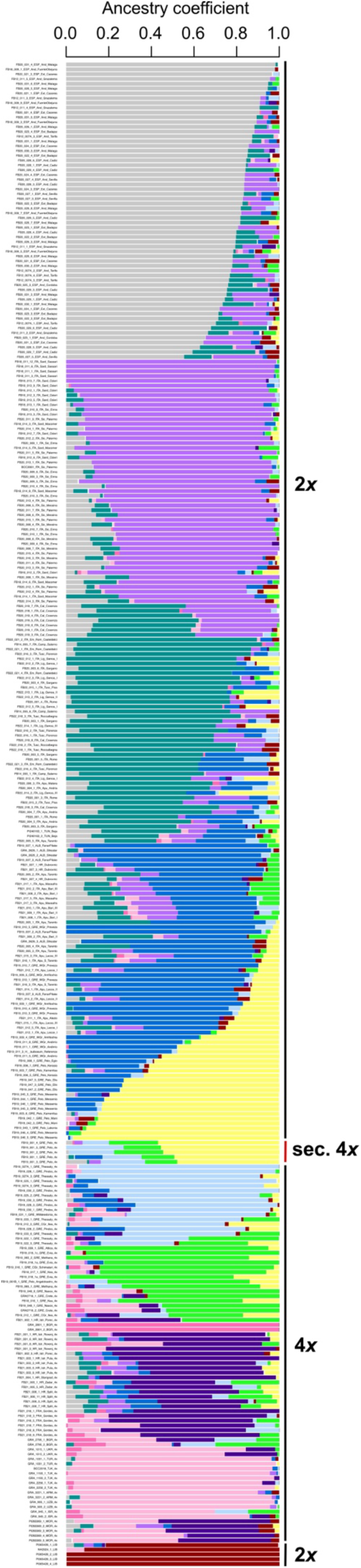
Bayesian population assignment plot of diploid and tetraploid cytotypes of *H. bulbosum* derived from an analysis in LEA at K = 11.

**Figure S7.**
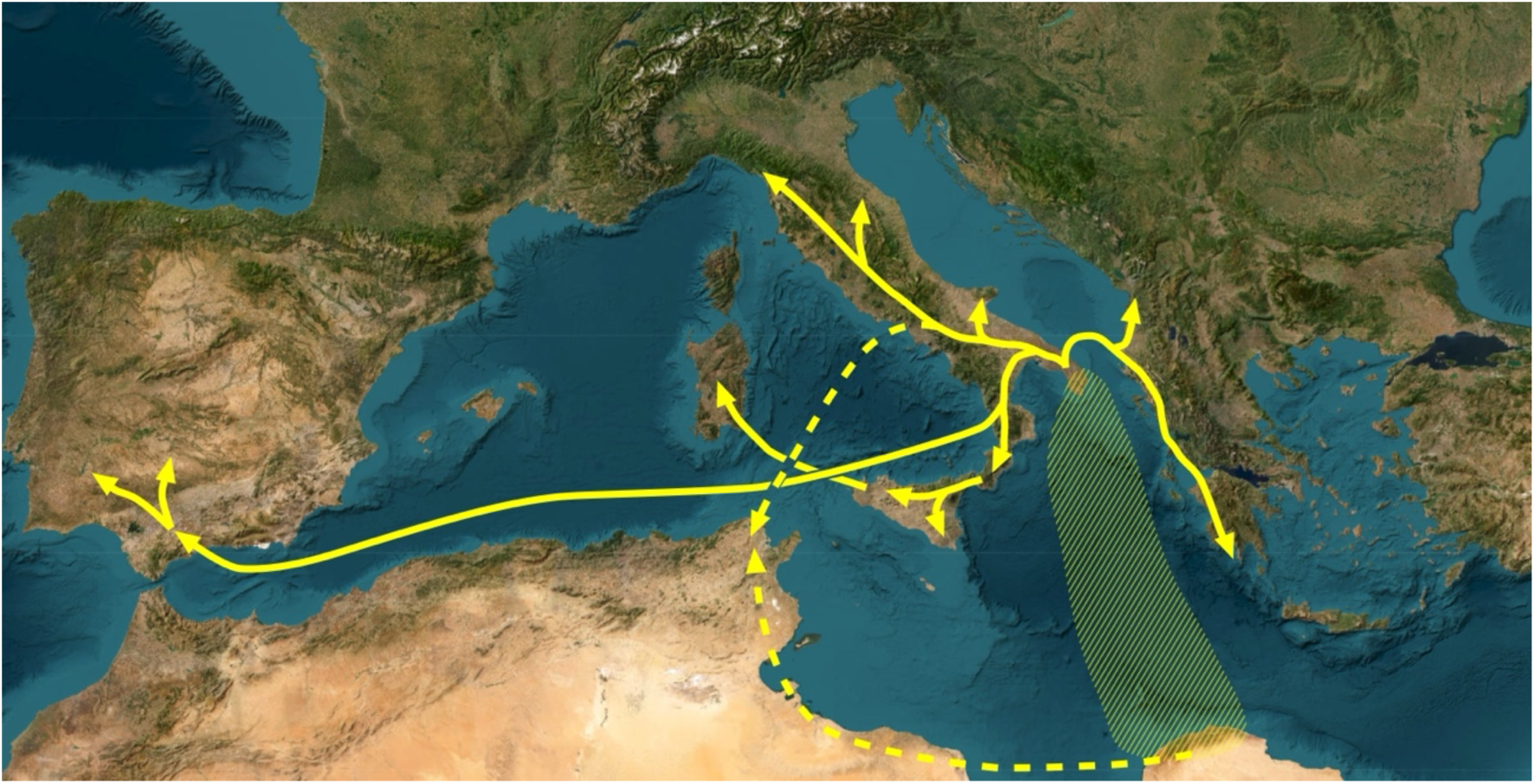
Colonization routes for the spread of diploid *H. bulbosum* through the central and western parts of the Mediterranean basin based on phylogeographic analyses of GBS data. The hatched area indicates that the region of origin of the species is currently not known but should be in northern Africa and/or southern Italy or more east of these regions. For the European part of the Mediterranean, we postulate an “out of Apulia” scenario with eastward colonization into Albania and Greece, a northward extension into mainland Italy, and westward colonization of Sardinia through Calabria and Sicily. Calabria was also the starting point for long-distance dispersal to the Iberian Peninsula. Tunisia was according to MP analyses reached by long-distance dispersal from mainland Italy, while an SVDQ analysis could imply a North African origin. The underlying map of the Mediterranean was created in Free-map.org (https://free-map.org).

## References

1. Ala N, Bagheri A, Zare H, Himmelbach A, Harpke D, Blattner FR. 2025. Merging GBS datasets to analyze the phylogeny of western Eurasian lime trees (*Tilia*) and place the Hyrcanian Forest taxa. BMC Plant Biology xx

2. Blattner FR. 2004. Phylogenetic analysis of *Hordeum* (Poaceae) as inferred by nuclear rDNA ITS sequences. Molecular Phylogenetics and Evolution 33: 289–299. 10.1016/j.ympev.2004.05.012

3. Blattner FR. 2006. Multiple intercontinental dispersals shaped the distribution area of *Hordeum* (Poaceae). New Phytologist 169: 603–614. 10.1111/j.1469-8137.2005.01610.x

4. Blattner FR. 2009. Progress in phylogenetic analysis and a new infrageneric classification of the barley genus *Hordeum* (Poaceae:Triticeae). Breeding Science 59: 471–480. 10.1270/jsbbs.59.471

5. Blattner FR. 2018. Taxonomy of the genus Hordeum and barley (Hordeum vulgare). In: Stein N, Muehlbauer GJ, eds. The barley genome. Cham: Springer, 11–23. 10.1007/978-3-319-92528-8_2

6. Bothmer R von, Jacobsen N, Baden C, Jorgensen RB, Linde-Laursen I. 1995. An ecogeographical study of the genus Hordeum (2nd edition). Rome: IPGRI.

7. Brassac J, Blattner FR. 2015. Species-level phylogeny and polyploid relationships in *Hordeum* (Poaceae) inferred by next-generation sequencing and in-silico cloning of multiple nuclear loci. Systematic Biology 64: 792– 808. 10.1093/sysbio/syv035

8. Cauderon Y, Cauderon A. 1956. Etudes de l’hybride F1 entre *Hordeum bulbosum* L. et *H. secalinum* Schreb. Annales de l’amélioration des plantes 6: 307–317.

9. Chapman EA, Thomsen HC, Tulloch S, Correia PMP, Luo G, Najafi J, DeHaan LR, Crews TE, Olsson L, Lundquist PO et al. 2022. Perennials as future grain crops: Opportunities and challenges. Frontiers in Plant Science 13: 898769. 10.3389/fpls.2022.898769

10. Danecek P, Auton A, Abecasis G, Albers CA, Banks E, DePristo MA, Handsaker RE, Lunter G, Marth GT, Sherry ST et al. 2011. The variant call format and VCFtools. Bioinformatics 27: 2156–2158. 10.1093/bioinformatics/btr330

11. Earl DA, von Holdt BM. 2012. STRUCTURE HARVESTER: A website and program for visualizing STRUCTURE output and implementing the Evanno method. Conservation of Genetic Resources 4: 359–361. 10.1007/s12686-011-9548-7

12. Eaton DAR, Overcast I. 2020. ipyrad: Interactive assembly and analysis of RADseq datasets. Bioinformatics 36: 2592–2594. 10.1093/bioinformatics/btz966

13. El Rabiai GT, Al tira FM, Lamlom SH. 2010. Preliminary checklist for the flora of Wadi El Ghattara in Libya. *Journal of Science and its Applications* (Garyounis University, Libya) 4: 39–47.

14. Elshire RJ, Glaubitz JC, Sun Q, Poland JA, Kawamoto K, Buckler ES, Mitchell SE. 2011. A robust, simple genotyping-by-sequencing (GBS) approach for high diversity species. PLoS ONE 6: e19379. 10.1371/journal.pone.0019379

15. Feng JW, Pidon H, Cuacos M, Himmelbach A, Fuchs J, Haberer G, Lux T, Kuo YT, Guo Y, Jayakodi M, et al. 2025. A haplotype-resolved pangenome of the barley wild relative *Hordeum bulbosum*. Research Square: 10.21203/rs.3.rs-3916840/v1

16. Fick SE, Hijmans RJ. 2017. WorldClim 2: New 1-km spatial resolution climate surfaces for global land areas. International Journal of Climatology 37: 4302–4315. 10.1002/joc.5086

17. Frichot E, François O. 2015. LEA: An R package for landscape and ecological association studies. Methods in Ecology and Evolution 6: 925–929. 10.1111/2041-210x.12382

18. Fuerst D, Shermeister B, Mandel T, Hübner S. 2023. Evolutionary conservation and transcriptome analyses attribute perenniality and flowering to day-length responsive genes in bulbous barley (*Hordeum bulbosum*). Genome Biology and Evolution 15: evac168. 10.1093/gbe/evac168

19. Hijmans R. 2023. raster: Geographic data analysis and modeling. R package version 3.6-26. https://CRAN.R-project.org/package=raster (accessed on 07 January 2023).

20. Jakob SS, Meister A, Blattner FR. 2006. The considerable genome size variation of *Hordeum* species (Poaceae) is linked to phylogeny, life form, ecology, and speciation rates. Molecular Biology and Evolution 21: 860– 869. 10.1093/molbev/msh092

21. Jakobsson M, Rosenberg NA. 2007. CLUMPP: A cluster matching and permutation program for dealing with label switching and multimodality in analysis of population structure. Bioinformatics 23: 1801–1806. 10.1093/bioinformatics/btm233

22. Jensen CJ. 1977. Monoploid production by chromosome elimination. In: Reinert J, Bajaj YPS, eds. Plant cell, tissue and organ culture. Berlin, Springer: 299–340.

23. Johnston PA, Timmerman-Vaughan GM, Farnden KJF, Pickering R. 2009. Marker development and characterisation of *Hordeum bulbosum* introgression lines: A resource for barley improvement. Theoretical and Applied Genetics 118: 1429–1437. 10.1007/s00122-009-0992-7

24. Jørgensen RB. 1982. Biosystematics of *Hordeum bulbosum* L. Nordic Journal of Botany 2: 421–434. 10.1111/j.1756-1051.1982.tb01205.x

25. Katznelson J, Zohary D. 1967. Diploid and tetraploid *Hordeum bulbosum* L. Israel Journal of Botany 16: 57–62.

26. Konzak CF, Randolph LF, Jensen NF. 1951. Embryo culture of barley species hybrids: Cytological studies of *Hordeum sativum* × *Hordeum bulbosum*. Journal of Heredity 42: 125–134. 10.1093/oxfordjournals.jhered.a106182

27. Lemon J. 2006. Plotrix: A package in the red light district of R. R-News 6: 8–12. https://journal.r-project.org/articles/RN-2006-026

28. Levin DA. 1975. Minority cytotype exclusion in local plant populations. Taxon 24: 35–43. 10.2307/1218997

29. Lischer HEL, Excoffier L. 2012. PGDSpider: An automated data conversion tool for connecting population genetics and genomics programs. Bioinformatics 28: 298–299. 10.1093/bioinformatics/btr642

30. Martin M. 2011. Cutadapt removes adapter sequences from high-throughput sequencing reads. EMBnet.journal 17: 10–12. 10.14806/ej.17.1.200

31. Minh BQ, Schmidt HA, Chernomor O, Schrempf D, Woodhams MD, von Haeseler A, Lanfear R. 2020. IQ-TREE 2: New models and efficient methods for phylogenetic inference in the genomic era. Molecular Biology and Evolution 37: 1530–1534. 10.1093/molbev/msaa015

32. Muscarella R, Galante PJ, Soley-Guardia M, Boria RA, Kass HM, Uriarte M, Anderson RP. 2014. ENMeval: An R package for conducting spatially independent evaluations and estimating optimal model complexity for Maxent ecological niche models. Methods in Ecology and Evolution 5: 1198–1205. 10.1111/2041-210X.12261

33. Ortiz L. A. González A, Chueca MC. 1985. On the presence of diploid and tetraploid forms of *Hordeum bulbosum* L. in Spain. Anales Jardín Botánico de Madrid 41: 361–365.

34. Papeš D, Bosiljevac V. 1984. Tetraploid populations of bulbous barley (*Hordeum bulbosum* L.). Acta Botanica Croatica 43: 335–340. https://hrcak.srce.hr/index.php/159049

35. Phillips SJ, Anderson RP, Schapire RE. 2006. Maximum entropy modeling of species geographic distributions. Ecological Modelling 190: 231–259. 10.1016/j.ecolmodel.2005.03.026

36. Pohler W, Szigat G. 1982. Versuche zur rekombinativen Genübertragung von der Wildgerste *Hordeum bulbosum* auf die Kulturgerste *H. vulgare*. 1. Mitt. Die Rückkreuzung VV × BBVV. Archiv für Züchtungsforschung 12: 87–100.

37. Pritchard JK, Stephens M, Donnelly P. 2000. Inference of population structure using multilocus genotype data. Genetics 155: 945–959. 10.1093/genetics/155.2.945

38. Schoener TW. 1968. *Anolis* lizards of Bimini: Resource partitioning in a complex fauna. Ecology 49: 704–726. 10.2307/1935534

39. Sherif AS, Mahklouf MH. 2020. An exclusive revision and annotated catalogue to the Grass Family of Libya. Species 21: 1–31. https://www.discoveryjournals.org/Species/current_issue/2020/v21/n67/A1.pdf

40. Soltis DE, Soltis PS. 1992. Molecular data and the dynamic nature of polyploidy. Critical Reviews in Plant Sciences 12: 243–273. 10.1080/07352689309701903

41. Soltis DE, Soltis PS. 1999. Polyploidy: Recurrent formation and genome evolution. Trends in Ecology and Evolution 14: 348–352. 10.1016/S0169-5347(99)01638-9

42. Subrahmanyam NC, Bothmer R von. 1987. Interspecific hybridization with *Hordeum bulbosum* and development of hybrids and haploids. Hereditas 106: 119–127. 10.1111/j.1601-5223.1987.tb00244.x

43. Swofford DL. 2002. PAUP*. Phylogenetic Analysis Using Parsimony (*and other methods). Version 4.0a169. Sunderland: Sinauer Associates.

44. Walther U, Rapke H, Proeseler G, Szigat G. 2000. *Hordeum bulbosum*–a new source of disease resistance– transfer of resistance to leaf rust and mosaic viruses from *H. bulbosum* into winter barley. Plant Breeding 119: 215–218. 10.1046/j.1439-0523.2000.00475.x

45. Warren DL, Glor RE, Turelli M. 2008. Environmental niche equivalency versus conservatism: Quantitative approaches to niche evolution. Evolution 62: 2868–2883. 10.1111/j.1558-5646.2008.00482.x

46. Wendler N, Mascher M, Himmelbach A, Johnston P, Pickering R, Stein N. 2015. Bulbosum to go: A toolbox to utilize *Hordeum vulgare*/*bulbosum* introgressions for breeding and beyond. Molecular Plant 8: 1507–1519. 10.1016/j.molp.2015.05.004.

47. Wendler N, Mascher M, Nöh C, Himmelbach A, Scholz U, Ruge-Wehling B, Stein N. 2014. Unlocking the secondary gene-pool of barley with next-generation sequencing. Plant Biotechnology Journal 12: 1122– 1131. 10.1111/pbi.12219

48. Westerbergh A, Lerceteau-Köhler E, Sameri M, Bedada G, Lundquist PO. 2018. Toward the development of perennial barley for cold temperate climates—Evaluation of wild barley relatives as genetic resources. Sustainability 10: 1969. 10.3390/su10061969

49. Wickham H. 2016. ggplot2: Elegant graphics for data analysis. New York, Springer: 1–189. 10.1007/978-0-387-98141-3

50. Wickham H, Wickham MH. 2022. Package ‘Tidyverse*’*. 2019: 1–5. Available online: http://tidyverse.tidyverse.org (accessed on 20 July 2022).

